# Construction of a multi-tissue cell atlas reveals cell-type-specific regulation of molecular and complex phenotypes in pigs

**DOI:** 10.1101/2023.06.12.544530

**Authors:** Lijuan Chen, Houcheng Li, Jinyan Teng, Zhen Wang, Xiaolu Qu, Zhe Chen, Xiaodian Cai, Haonan Zeng, Zhonghao Bai, Jinghui Li, Xiangchun Pan, Leyan Yan, Fei Wang, Lin Lin, Yonglun Luo, Goutam Sahana, Mogens Sandø Lund, Maria Ballester, Daniel Crespo-Piazuelo, Peter Karlskov-Mortensen, Merete Fredholm, Alex Clop, Marcel Amills, Crystal Loving, Christopher K. Tuggle, Ole Madsen, Jiaqi Li, Zhe Zhang, George E. Liu, Jicai Jiang, Lingzhao Fang, Guoqiang Yi

## Abstract

The systematic characterization of cellular heterogeneity among tissues and cell-type-specific regulation underlying complex phenotypes remains elusive in pigs. Within the Pig Genotype-Tissue Expression (PigGTEx) project, we present a single-cell transcriptome atlas of adult pigs encompassing 229,268 high-quality nuclei from 19 tissues, annotated to 67 major cell types. Besides cellular heterogeneity within and across tissues, we further characterize prominent tissue-specific features and functions of muscle, epithelial, and immune cells. Through deconvoluting 3,921 bulk RNA-seq samples from 17 matching tissues, we dissect thousands of genetic variants with cell-type interaction effects on gene expression (ieQTL). By colocalizing these ieQTL with variants associated with 268 complex traits, we provide new insights into the cellular mechanisms behind these traits. Moreover, we highlight that orthologous genes with cell-type-specific regulation in pigs exhibit significant heritability enrichment for some human complex phenotypes. Altogether, our work provides a valuable resource and highlights novel insights in cellular regulation of complex traits for accelerating pig precision breeding and human biomedical research.

## Introduction

The cell is a fundamental structural, biological, and evolutionary unit of life and plays a key role in orchestrating the development and homeostasis of all living beings through global intercellular interactions. Multicellular organisms, including mammals, are generally composed of over 400 distinct cell types that are distinct in morphology and function (*1–5*). Genome-wide association studies (GWASs) have revealed that over 90% of phenotype-associated genetic variants lie within non-coding regions, suggesting that these variants might influence complex phenotypes through gene expression modulation (*6–8*). The limited overlaps between bulk expression quantitative trait loci (eQTL) and GWAS signals suggest that many candidate variants might regulate biological processes and then complex phenotypes through cell-type-specific mechanisms (*9–12*). Single-cell omics studies have shown that the substantial disorders in cellular activity, identity, and composition play a crucial role in the development of complex traits and diseases, both within and across individuals (*5, 13–16*), highlighting the importance of constructing a multi-tissue single-cell atlas for functionally understanding genotype-phenotype associations. In addition, a better understanding of molecular and cellular mechanisms underpinning complex phenotypes will be an important initial step in generating new avenues for precision breeding in agriculture and therapeutic solutions for similar human diseases (*13, 14, 17*).

As an important farm animal species, the domestic pig (*Sus scrofa*) is not only an abundant source of animal protein worldwide but also serves as a valuable human biomedical model and an optimal organ donor for xenotransplantation (*18*). Numerous studies in pigs have delineated significant QTL underlying complex traits of economic importance (*19, 20*), leading to vast improvements in pig breeding programs and production efficiency. However, the systematic interpretation of molecular mechanisms underlying complex phenotypes in pigs lags behind human and mouse research due to limitations in functional data availability. The ongoing Functional Annotation of Animal Genomes (FAANG) and Farm animal Genotype-Tissue Expression projects (FarmGTEx) are global efforts to provide catalogues of functional elements and variants in pigs at tissue level (*21–23*). The next step is to explore the cell-type-dependent biological consequences of trait-associated variants as tissues contain numerous cell types (*24*). Although some studies have conducted single-cell/nucleus RNA-seq (scRNA-seq and snRNA-seq) analyses in pigs, they primarily focused on elucidating the cellular heterogeneity and trajectories of lineage specification in a limited range of tissue types (*25–32*). The cell-type-specific biological impacts of genetic variants on complex traits by integrating single-cell RNA-sequencing with large-scale pig genetics data still need to be explored.

To further fine-map the causative genetic variants and decipher their cellular impacts on both molecular and complex phenotypes in pigs, we first constructed a single-nucleus transcriptome atlas by profiling a total of 319,433 nuclei from 19 major tissue types, representing 261 major cell clusters. Dissection of muscle, epithelial and immune cells depicted the cellular heterogeneity across these tissues and revealed a number of critical master regulators (*i.e.*, GATA4 and ZBTB11) driving cell identity. Through cellular deconvolution of PigGTEx tissues, cell-type interaction expression QTL (ieQTL) mapping, and the integrative analysis with GWAS results of 268 complex traits, we pinpoint the cell-type-specific contexts in which trait-associated genetic variants regulate the transcriptional activity and result in phenotypic variation. Moreover, we demonstrate that orthologous genes with cell-type-specific regulation in pigs exhibit significant heritability enrichment for many human complex phenotypes. Overall, this study enriches and enhances rich and open resources (http://piggtex.farmgtex.org/ and https://dreamapp.biomed.au.dk/pigatlas/) for charting the cell-cell transcriptome variability within and across tissues and expands our understanding of the connections between genetic variants and phenotypes at single-cell resolution in pigs. Our results provide relevant information for the development of future precision breeding strategies in pigs and human biomedical research.

## Results

### Global landscape of single-nucleus transcriptomic reference atlas from 19 pig tissues

To generate a comprehensive multi-tissue single-cell transcriptomic reference atlas of pigs, we performed snRNA-seq in 19 tissues/organs from two adult Meishan pigs (one male and one female) using 10 × Genomics technology, including subcutaneous adipose, cerebellum, cerebrum, colon, duodenum, heart, hypothalamus, ileum, jejunum, kidney, liver, lymph node, skeletal muscle, ovary, pancreas, pituitary gland, spleen, testis, and uterus (Fig. 1a). Initially, we profiled 16,812 nuclei and sequenced over 660 million raw reads per tissue on average (Fig. 1a). After quality control (see Methods for details), we obtained transcriptomic data for a total of 229,268 high-quality nuclei across all the 19 tissues (Supplementary Fig. 1). We first assessed the transcriptional similarity by comparing our snRNA-seq data with that from a previous study across seven common tissues (*27*). The Spearman correlation values between the two pseudo-bulk single-cell transcriptomic profiles were high for all tissues, ranging from 0.653 to 0.825 (Supplementary Fig. 2), suggesting globally consistent transcriptional profiles of samples between these two single-cell RNA-seq studies at the bulk tissue level.

**Fig. 1.**
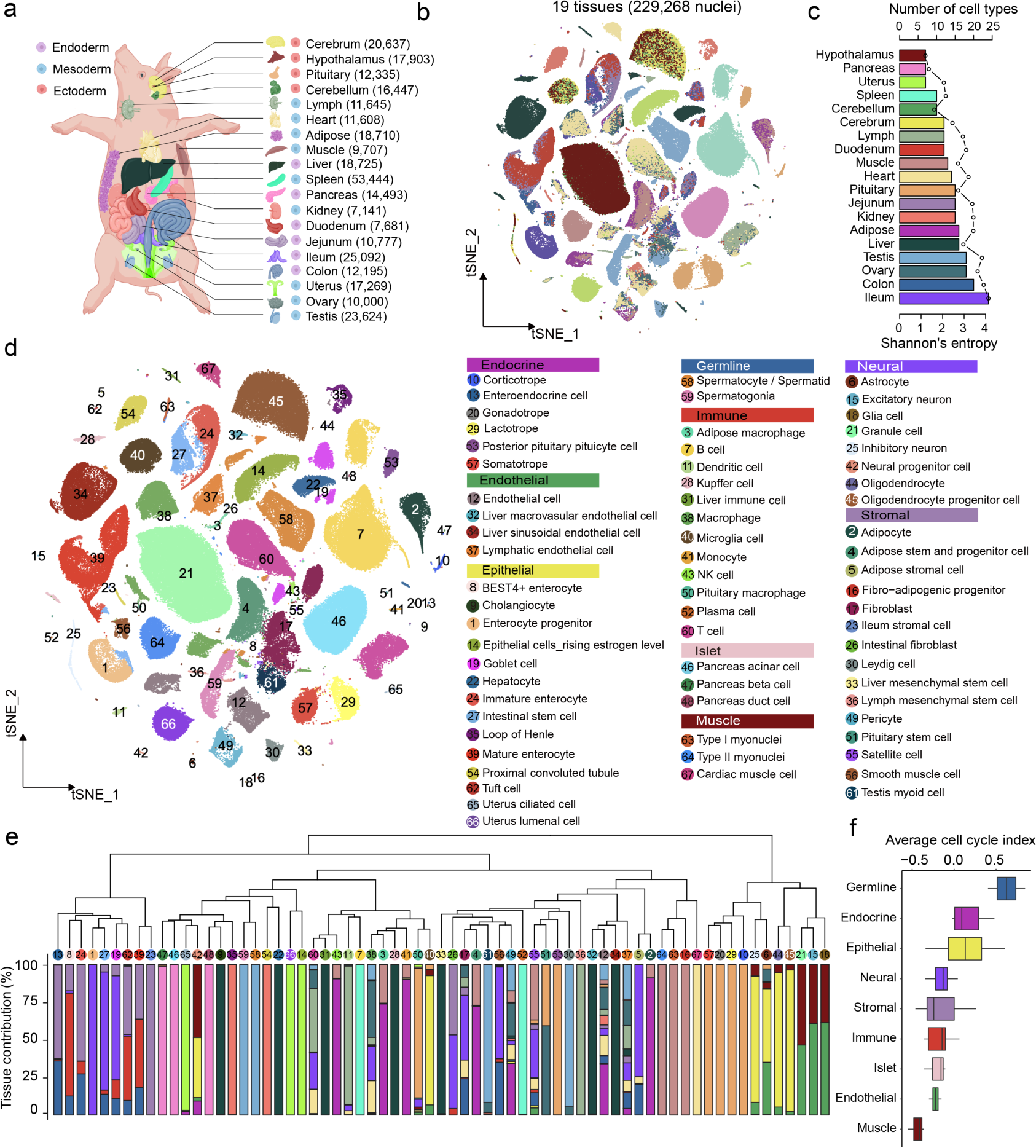
Single-nucleus transcriptomic landscape across 19 frozen tissues in adult pigs. **a,** Schematic diagram showing 19 primary pig tissues collected for snRNA-seq in this study. The cartoons used to generate this illustration were purchased from BioRender.com. The number of nuclei profiled per tissue is denoted in parentheses. **b,** t-SNE visualization of single-nucleus profiles (dots) colored by tissues. **c,** Bar plot displaying the number and diversity of cell types identified in each of the 19 tissues. Entropy shown by dotted line was calculated as described in Methods. **d,** t-SNE visualization of single-nucleus profiles (dots) colored by major cell types. All cell types are categorized into nine top-level cell lineages, and cell type annotation is provided in the legend to the right. **e,** Cellular relationship and composition across tissues. The dendrogram was created by hierarchical clustering based on the transcriptional levels of each cell type. The bar chart represents the relative contributions of tissues to each cell type. **f,** Cell state prediction of nine top-level cell lineages. Cells with higher cell cycling index are more proliferative. The horizontal line in the boxplots corresponds to the median, the box bounds indicate the 25th and 75th percentiles and the whiskers represent 1.5 times the interquartile range. Values outside the whiskers are displayed as points.

The complete snRNA-seq dataset was grouped into 77 cell clusters and manually annotated as 67 major cell types based on the expression of canonical marker genes from the literature (Fig. 1b-c, Supplementary Fig. 3, and Supplementary Table 1). All tissues and cell types showed sufficient transcriptional abundance, with a median of 4,008 unique molecular identifiers (UMI) and 2,064 transcribed genes per nucleus, therefore displaying higher expression than the previously reported single-cell data in pigs (*27*). The global cell atlas revealed that a majority of cell types, like cardiomyocytes, enterocytes, and hepatocytes, exhibited a high tissue specificity regarding gene expression (Fig. 1d), reflecting their specialized functions. Notably, several prevalent cell types, such as immune cells, endothelial cells, and fibroblasts, were commonly shared among tissues. To gain a deeper understanding of cellular heterogeneity within each tissue, we generated individual visualizations in the hierarchy with Uniform Manifold Approximation and Projection (UMAP), resulting in an average of 14 main cell types per tissue (Supplementary Figs. 4-5). Of note, the ileum showed 24 putative cell subpopulations, consistent with its highest cell-type diversity evaluated by the Shannon entropy index (Fig. 1c-d and Supplementary Figs. 4-5). Additionally, we compared cellular signatures of tissues shared by our work and the previous study (*27*), and in general, found a high consistency in both cell annotation, distribution, and expression (Supplementary Fig. 6). However, some cell types or marker genes, such as *ADIPOQ* in adipose tissue, *DOCK4* in heart, and *CD163* in liver, displayed distinct expression levels and patterns between the two studies (Supplementary Fig. 6). This discrepancy might be attributed to differences in tissue sampling regions and experimental protocols. To further probe the intercellular relationships, we conducted an unsupervised hierarchical clustering analysis for all these 67 cell types based on their transcriptomic profiles (Fig. 1e). These cell types could be largely classified into nine different functional groups of cells, including endocrine, endothelial, epithelial, germline, immune, islet, muscle, neural and stromal cells. Remarkably, we observed a higher similarity among cell types within the nine major lineages rather than among tissues, suggesting that cell clustering was primarily driven by cell type and that these neighboring cell types possibly had similar functions (Fig. 1e). To evaluate the dynamics of cell state in each cell type, we computed the cell cycling index as described previously (*3*). Germline cells exhibited a greater cell division capacity than other cells, while the endothelial, stromal, and muscle compartments, which are known to be largely quiescent, had low cycling indices (Fig. 1f and Supplementary Fig. 7). Notably, the epithelial cells presented the highest variations in cell states, suggesting great functional diversity of epithelial cell subpopulations.

### Distinct transcriptional patterns among three types of muscle cells

The muscular system is a complex collection of organs that allow movement through the contraction of muscle fibers and is also the main production target of the pig industry, with the aim to provide high-quality protein in the form of meat. There are three distinct muscle types in the body, namely skeletal, cardiac, and smooth muscle, each with unique cellular morphologies and functions (*33*). We found that skeletal muscle cells and cardiomyocytes accounted for 66.43% and 25.09% of total cells in muscle and heart, respectively, while smooth muscle cells could be found in eight tissues with an average proportion of 2.52% (Supplementary Figs. 4-5). To provide a more detailed view of the three muscle cell types, we extracted a total of 10,117 muscle cells from corresponding tissues according to cell type annotations and performed the dimension reduction analysis. As expected, t-SNE inspection and dendrogram showed a clear separation among the three major muscle cell types, and each specifically expressed its classical marker genes (Fig. 2a-b), like *MYH7*, *MYBPC2,* and *TNNT1* for skeletal muscle cells, *MYBPC3* and *TNNT2* for cardiac muscle cells, and *ACTA2*, *MYH11*, and *RYR2* for smooth muscle cells. We observed a preferential grouping of skeletal muscle cells with cardiac muscle cells since both belong to striated muscle tissue and share similar structural and functional characteristics (*34*). In addition, skeletal and smooth muscle cells could be further partitioned into multiple subclusters in the hierarchy (Fig. 2b), suggesting their subtle context-dependent functions. To examine global transcriptional differences among the three muscle cell types, we performed a pair-wise differential gene expression analysis. In total, we identified 1,250 differentially expressed genes (DEGs) across the three myocyte subtypes (Fig. 2c). The 343 DEGs in skeletal muscle cells were significantly enriched in striated muscle contraction, while DEGs in cardiac muscle cells were mainly involved in cardiac muscle tissue development. Smooth muscle cell-specific genes were enriched in the extracellular matrix organization (Fig. 2c). Further analysis of transcription factor (TF) activity revealed many remarkable regulons in the control of muscle cell type specification (Fig. 2d). For example, MYOD1, MYOG, and FOXO4 served as master TFs responsible for skeletal muscle cell development, while certain members of the GATA and TBX families showed unique regulatory roles in the cardiac and smooth muscle cell types, respectively.

**Fig. 2.**
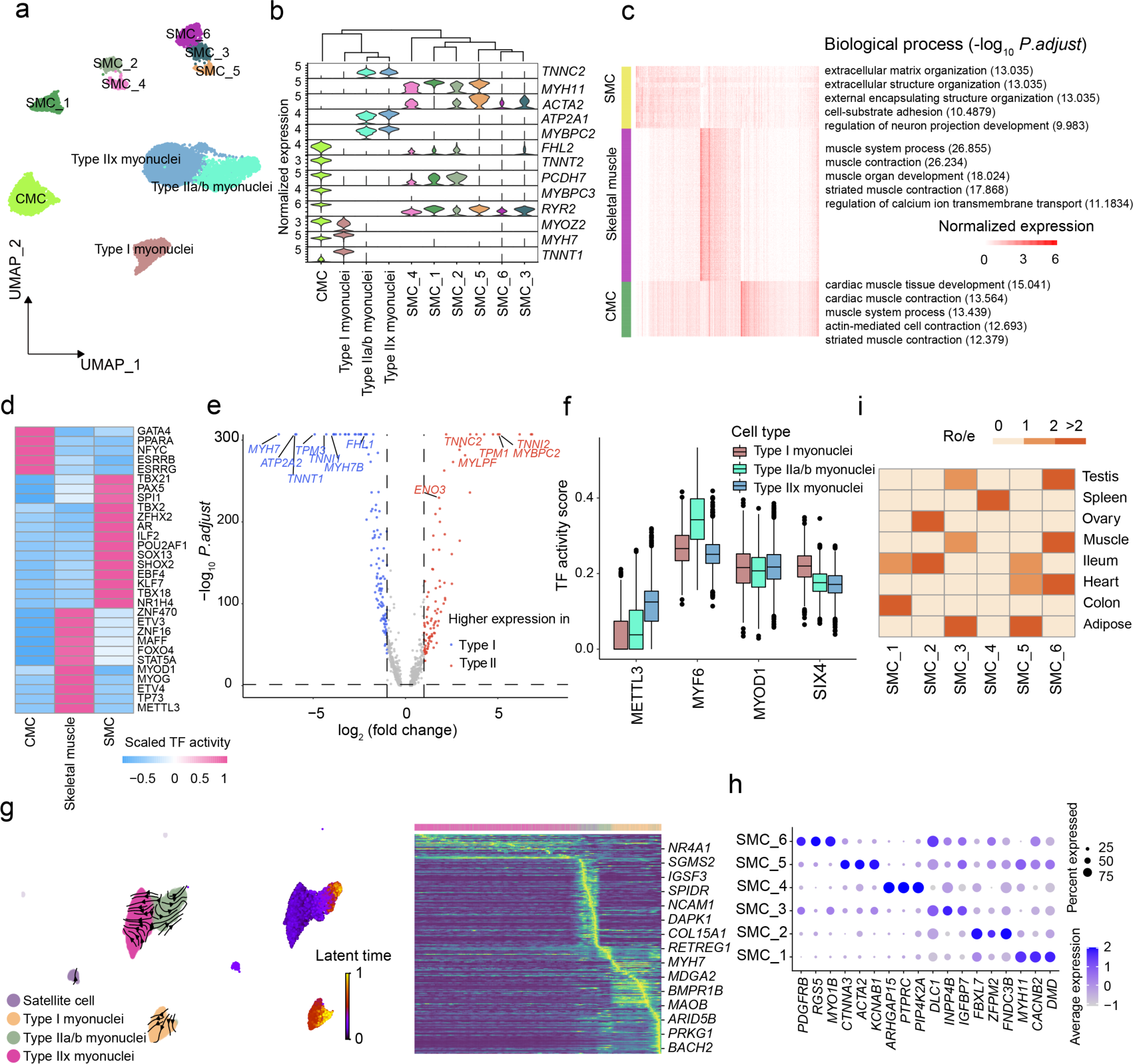
Identification and characterization of three muscle cell types. **a,** UMAP visualization of all muscle cells from eight tissues. Each dot represents one nucleus, with colors coded according to manually annotated cell types. CMC, cardiac muscle cell; SMC, smooth muscle cell. **b,** Violin plots showing the normalized expression levels of marker genes for three major muscle cell types. **c,** Significantly enriched biological process terms of specific gene signatures in three major muscle cell types. Numbers between parentheses represent significance expressed as −log_10_ (adjusted *p*-value). **d,** Transcription factors with different activity scores among three major muscle cell types. **e,** Volcano plot displaying differentially expressed genes between type I and type II myonuclei. **f,** Four candidate transcription factors with distinct activity scores in type I, IIa/b, and IIx myonuclei. **g,** RNA velocity analysis demonstrating state transition from satellite cells to myofiber in the skeletal muscle tissue. The arrows represent a flow derived from the ratio of unspliced to spliced transcripts, which in turn predicts dynamic changes in cell identity. Heatmap on the right demonstrates stereotyped changes in gene expression trajectory. **h,** Dot plot showing the expression levels of selected marker genes for each smooth cell cluster. **i,** Heatmap indicating the tissue preference of each cell population across different tissues revealed by R_o/e_ (ratio of observed cell number to expected cell number).

In addition to characterizing differences across the three main muscle cell types, we further probed cellular heterogeneity within skeletal and smooth muscle cells separately. Our analysis of pig myonuclei in skeletal muscle confirmed the presence of *MYH7* type I (slow-twitch) and *TNNC2* type II (fast-twitch, IIa/b, and IIx) myofibers (Fig. 2b), consistent with a previous study in monkeys (*5*). A pairwise comparison between type I and type II myofibers uncovered 209 DEGs (Fig. 2e). Notably, type I myofiber-specific genes were enriched in several fundamental pathways related to molecular structure and function like muscle contraction and sarcomere organization (Supplementary Fig. 8a), while the upregulated genes in type II myofibers were essential for metabolic pathways such as phosphorylation and glycolysis (Supplementary Fig. 8b). By examining DEGs of these two types of myofibers previously reported in humans (*35, 36*), we observed a strong positive Pearson correlation of 0.945 regarding fold changes of the shared genes between pigs and humans (Supplementary Fig. 8c), implying that the process of muscle fiber specialization might be highly conserved between these two species. Furthermore, we identified several critical master regulators, including METTL3, MYF6, and SIX4, which displayed distinct regulatory activities in type I, IIa/b, and IIx myonuclei (Fig. 2f). By conducting RNA velocity analysis in myofibers together with satellite cells (known as skeletal muscle stem cells), we further explored the differentiation trajectory of muscle fibers. Our results revealed clear myogenesis from satellite cells to mature muscle fibers (Fig. 2g), which were driven by several fundamental genes with dynamic expressions across distinct cell states such as *MYH7* and *PRKG1*. Interestingly, the type IIa/b fibers displayed intermediate cell states and characteristics during the fast-to-slow fiber-type switch. In the smooth muscle cell compartment, we found distinct gene signatures and tissue enrichment among these six cell subtypes (Fig. 2h-i). For instance, SMC_1, which was preferably located in the intestine, showed much higher activity of *MYH11* and *DMD*, while SMC_6, mainly from testis, exhibited exclusively high expression of *MYO1B* and *RGS5*. These results suggested that the same cell types undergo subtle processes of functional differentiation depending on the original tissue contexts in which they reside.

### About the similarity and heterogeneity of intestinal epithelial cells

Epithelia are sheets of cells that cover most body surfaces, line internal cavities, and compose certain glands. They perform a wide range of biological functions, including protection, absorption, and secretion (*37*). First, we pursued to investigate the primary characteristics and functions of epithelial cells, given their high abundance and diversity in the different organs. We obtained a total of 57,049 epithelial cells from eight tissues and identified their tissue-specific expression patterns and functions through the global t-SNE and hierarchical clustering (Fig. 3a and Supplementary Fig. 9). Epithelial cells from the duodenum, jejunum, ileum, and colon, representing the digestive system in the present study, exhibited closer relationships with other cells from the same digestive system than with cells from other systems. As expected, epithelial cells from the intestines showed a strong digestive and metabolic capacity, such as microvillus organization and intestinal absorption, compared to other subtypes (Fig. 3b-c). We then extracted intestinal stem cells, enterocytes, and enteroendocrine cells for further exploration, as these cell types might play pivotal roles in feed efficiency traits in pigs (*38–40*). Intestinal stem cells expressed high levels of *OLFM4* and *LGR5* and could be further subdivided into two subtle subtypes according to the differential expression levels of these two markers (Fig. 3d). We defined four enterocyte subgroups by the transcriptional patterns of canonical enterocyte markers (for example, *MUC13*, *SI*, *FUT8*, *APOB*, and *BEST4*), including enterocyte progenitors, immature enterocytes, mature enterocytes, and BEST4^+^ enterocytes. Enteroendocrine cells, which are specialized gut epithelial cells that produce and release hormones in the intestine (*40*), displayed a higher expression of *RAB3C*, *CHGA*, and *STXBP5L* when compared to other intestinal epithelial cells. Enrichment analyses of cell types across tissues revealed that intestinal stem cells were mainly located in the ileum and, to some extent, in the jejunum and colon, while enterocytes were more abundant in the duodenum and colon (Fig. 3e).

**Fig. 3.**
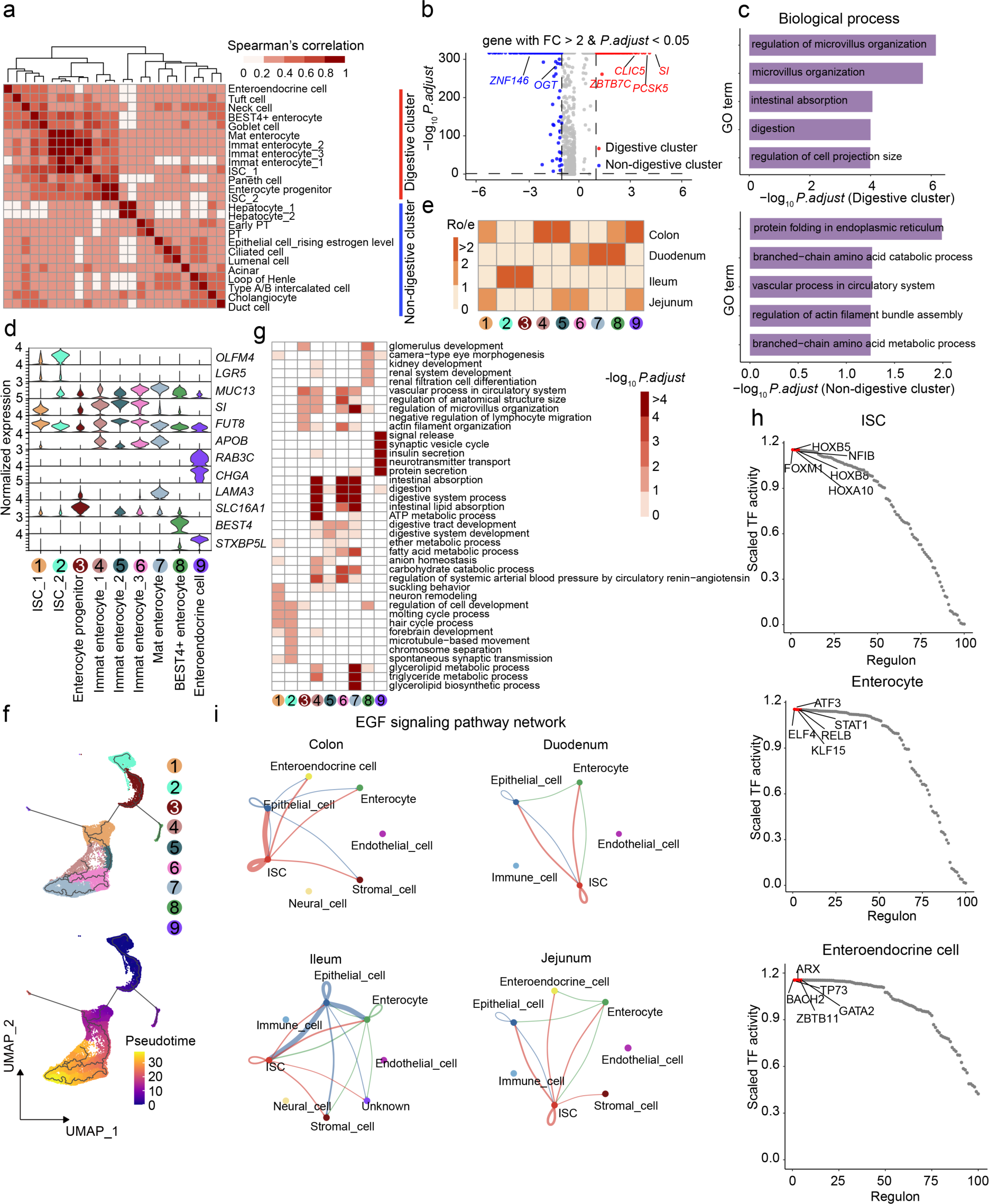
Shared and tissue-specific molecular features for epithelial cell compartments. **a,** Heatmap showing Spearman correlation coefficient between 25 epithelial cell subtypes which could be broadly classified into digestive and non-digestive groups. **b,** Volcano plot displaying differentially expressed genes between the digestive and non-digestive clusters. Dots in the volcano plot highlight up-regulated genes in each group. **c,** Functional annotation of up-regulated genes in each group. Top enriched biological processes terms are listed. **d,** Violin plots showing the normalized expression levels of marker genes for each cell subtype. **e,** Heatmap indicating the tissue preference of each cell population across four intestinal segments revealed by R_o/e_ (ratio of observed cell number to expected cell number). **f,** UMAPs showing the pseudotime differentiation trajectories of intestinal stem cells, enterocytes, and enteroendocrine cells, respectively. **g,** Heatmap representing the enrichment of biological process terms in epithelial cell subtypes. **h,** Scatter plots showing the top 100 regulons of the three major epithelial cell subtypes. Each regulon is ordered by activity score, and the top five regulons with high activity are highlighted in red. **i,** The inferred EGF signalling pathway network among the major cell types in four intestinal segments. The edge width represents the communication probability, and a thicker edge line indicates a stronger signal.

To further characterize the lineage relationships and cell states among intestinal stem cells, enterocytes, and enteroendocrine cells, we conducted the pseudotime analysis and cell cycling index prediction (*3, 41*). Both analyses revealed that intestinal stem cells and enterocyte progenitors exhibited a great capacity for differentiation into enterocytes and enteroendocrine cells, as evidenced by their high proliferative states (Fig. 3f and Supplementary Fig. 10a-b). The differentiation trajectory of these intestinal epithelial cells was highly similar among the four individual intestine segments (Supplementary Fig. 10c-f). Functional annotation analyses based on the Gene Ontology (GO) database demonstrated that gene signatures of each cell subgroup in intestinal stem cells were mainly enriched in cell cycle-related biological processes as expected (Fig. 3g). The highly expressed genes in BEST4^+^ enterocytes were over-represented in cell development and morphogenesis, which was highly different from the functions of immature and mature enterocytes. The gene sets restricted in enteroendocrine cells were significantly enriched in signal release and protein secretion (Fig. 3g). The distinct transcriptional profiles and functions of these cell types can be attributed to their diverse gene regulatory programs (Supplementary Fig. 11). By inferring the TF activity across the trajectory, we found that three master regulators, NFIB, STAT1, and ZBTB11, play essential roles in enterocyte lineage specification by a coordinated sequential activation (Fig. 3h and Supplementary Fig. 12). To compare the structures and intensities of cell-cell communication across the four gut segments, we employed CellChat (*42*) to identify potential ligand-receptor pairs among the major cell types. Our results revealed that EGF, PDGF, and BMP signaling pathways were major communicating pathways in the porcine intestine segments (Fig. 3i and Supplementary Fig. 13). Although the global interaction patterns were similar, the strength of intercellular interactions was different across intestine segments. For instance, compared with the colon, we observed stronger intercellular interactions among enterocytes, epithelial cells, and intestinal stem cells in small intestine tissues. We further mapped ligand-receptor pairs in specified cell subpopulations across different organs to understand the rewiring of molecular interactions regulating cell-cell interactions. Notably, the “NAMPT-INSR” and “GHRL-GHSR” ligand-receptor pairs were specific in interactions between enterocytes. Overall, our findings highlight the importance of dynamic information exchange between different cells in contributing to the diverse digestive functions of different intestine sections.

### A cross-tissue reference of immune cell types and states

The immune system is a complex network of cell types distributed throughout the whole body and provides protection against bacteria, viruses, and other pathogens. Understanding the specific and shared features of tissue-resident immune cells is crucial for deciphering the molecular mechanisms underlying immune responses and ultimately for accelerating precision breeding of disease resistance in pigs. We identified a total of 45,491 immune cells prevailing in 17 tissues, including T cells, B cells, natural killer cells (NK), macrophages, and other tissue-resident immune cells (Fig. 4a and Supplementary Fig. 14). Hierarchical clustering analysis revealed three main branches of immune cells: myeloid and lymphoid lineages, as well as microglia, which are brain-resident macrophages (Fig. 4b). As expected, each tissue has its own immune compartments, with specific immune cell compositions. For example, the four major parts of the brain exclusively contain microglia cells. A large population of B cells was evident in the spleen, whereas lymph nodes were enriched for multiple T cell types. We next subdivided and reanalyzed the immune dataset to explore further heterogeneity within macrophages and T cells, which were abundantly present across tissues. All tissue-resident macrophages, together with monocytes, were divided into 13 more granular subsets, which were supported by the expression of well-established marker genes (Fig. 4c). These macrophage subgroups exhibited clear tissue-type separation and preference, although certain subsets were shared by multiple tissues (Fig. 4b). For instance, the M1 macrophage subgroups were enriched in muscle and liver, while M2 macrophages were mainly located in ileum and ovary. To further dissect cell-type-specific transcriptional profiling, we performed pair-wise differential expression analyses and identified 2,903 genes with restricted expression in one or a few cell types (Supplementary Fig. 15a). Functional enrichment analysis evidenced the presence of overrepresented biological processes for each macrophage subtype, which recapitulated cell-type-specific functions regarding resident tissues and niches as well as putative cellular states (Fig. 4d-f and Supplementary Fig. 15b). Furthermore, cell-type-specific transcriptional programs were combinatorially controlled by several TFs with overlapping expression patterns. The regulons KLF3 and CEBPB were exclusively expressed in monocyte subsets and showed a gradual decrease in expression levels across the monocyte-to-macrophage differentiation trajectory (Fig. 4g).

**Fig. 4.**
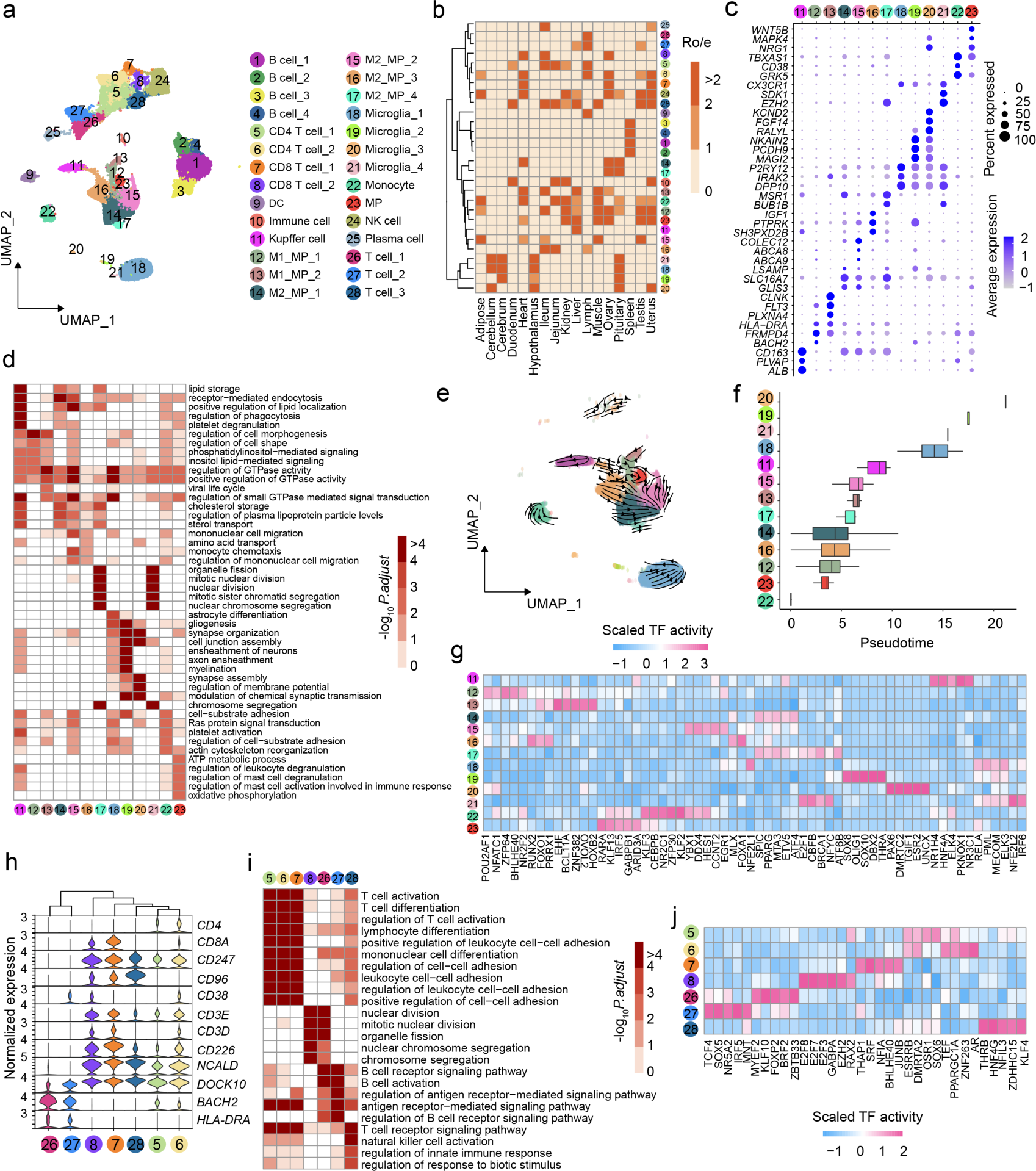
Immune cell heterogeneity across tissues in pigs. **a,** UMAP visualization of immune cell types across different tissues. Each dot represents one cell, with colors coded according to manually annotated cell types. **b,** Heatmap indicating the tissue preference of annotated immune cell types across different tissues revealed by R_o/e_ (ratio of observed cell number to expected cell number). **c,** Dot plot showing the expression levels of selected marker genes for each cell cluster. **d,** Heatmap representing the enrichment of biological process terms for monocyte and macrophage lineages residing in different tissues. **e,** UMAP showing the pseudotime differentiation trajectories of monocyte and macrophage lineages. **f,** Box plots denoting the distribution of estimated pesudotime value for each cell type by Monocle3. **g,** Heatmap showing transcription factors with distinct activity scores in six major myeloid cell compartments. **h,** Violin plots showing the normalized expression levels of marker genes for T cell populations. **i,** Heatmap representing the enrichment of biological process terms for T cell subtypes in different tissues. **j,** Heatmap showing transcription factors with different activity scores among different T cell subtypes.

T cells play a crucial role in elicitating and controlling the adaptive immune response (*43*). We identified seven T cell clusters based on known gene signatures, with CD4^+^ and CD8^+^ T cells showing a distinct separation, while the remaining clusters were designated as general T cells due to the absence of significant CD4 or CD8 surface molecules (Fig. 4h). CD4^+^ and CD8^+^ T cells in our data were further divided into two subtle clusters, respectively, based on the transcriptional differences of several classical markers like *CD3E* and *NCALD*. While these annotated T cell clusters were observed in 14 organs, their relative proportion and enrichment varied greatly across different organs (Fig. 4b). CD4^+^ T cells were primarily located in lymph nodes and jejunum, whereas CD8^+^ T and NK cells were more abundant in heart and ovary. To understand their potential diverse biological functions, we identified DEGs among these T cell subtypes and then carried out a functional annotation. The majority of T cells shared several enriched GO terms, like T cell activation and T cell receptor signaling pathway, suggesting their shared immune functions regardless of tissue origins. Specifically, signatures of CD4^+^ T cells were enriched for cell-cell adhesion, whereas CD8^+^ T cells had enhanced biological functions in nuclear division and regulation of antigen receptor-mediated signaling pathway (Fig. 4i and Supplementary Fig. 16). The distinct transcriptional profiles and molecular functions were attributed mainly to the specific TF network (Fig. 4j). Overall, our study provides valuable insights into the diversity and complexity of T cell populations across different organs and sheds light on their roles in regulating the immune response.

### Genetic mapping and functional implications of cell-type-specific eQTL

Bulk tissue samples often contain a high degree of cellular heterogeneity, which can mask genetic effects that are active only in specific cell types within the sampled tissue. To address this, we explored ieQTL by performing the cell-type deconvolution analysis of 3,921 bulk RNA-seq samples in the PigGTEx project via this newly built cross-tissue cell atlas. First, we tested the cell estimation performance of the CIBERSORT algorithm (*44*) in pigs by deconvoluting pseudo-bulk samples generated from simulation studies using the SCDC software (*45*). By employing the gene signature matrix built from our liver snRNA-seq data, we observed that the estimated cell proportions from pseudo-bulk samples were highly correlated with the putative cell populations identified in the liver snRNA-seq, with the highest correlation in Hepatocyte_1 subtype (Pearson’s *r* = 0.841, *p*-value < 2.2 ×10^-16^, Supplementary Fig. 17a-d). This result indicated the feasibility and accuracy of our cellular deconvolution pipeline in pigs. To identify cell-type-specific eQTL in an unbiased manner, we performed eQTL deconvolution analysis by integrating our cross-tissue snRNA-seq data with the large-scale bulk RNA-seq collections from the PigGTEx project.

The pseudo-bulk gene expressions of our snRNA-seq data were significantly correlated with those of PigGTEx bulk samples across all the 17 matching tissues, with correlation coefficients ranging from 0.498 (colon) to 0.745 (spleen), implying sufficient concordance for the subsequent integration (Supplementary Fig. 17e). We thus estimated the relative cell fractions of these 17 PigGTEx tissues using the snRNA-seq signature matrix of the respective tissues, where sample sizes of PigGTEx tissues varied from 44 (kidney) to 1,321 (muscle). Overall, most samples were well-deconvoluted (*p*-value < 0.05, 1,000-times permutations) and revealed a striking variability in cellular composition across the PigGTEx samples (Fig. 5a and Supplementary Fig. 18). The number of putative cell types detected in deconvoluted samples ranged from six (uterus) to 23 (ileum) (Supplementary Fig. 17f). In particular, the predicted abundance of cell types in muscle and heart displayed considerable inter-individual variations, with certain cell types in some samples even being totally missing, while colon and hypothalamus showed less heterogeneous cell fractions across samples (Supplementary Fig. 18). To map ieQTL, we performed a linear regression analysis that models an interaction term between estimated cell fractions and genotypes (*46*). We detected a total of 5,168 protein-coding genes with at least one significant ieQTL (ieGenes) across cell types and tissues (Fig. 5b), with around a third of these ieQTL validated using the allele-specific expression approach. Of note, muscle exhibited the highest number of significant ieGenes, followed by cerebrum, testis, liver, and adipose tissues (Fig. 5c). The discovery power of ieGenes in tissue was significantly correlated with its sample size (Fig. 5b). We detected an average of 114 ieGenes across 79 cell types from 14 tissues. Among them, type IIx myonuclei had the largest number of ieGenes (n = 797), whereas ileum Paneth cells only had one ieGene. For instance, the effects of *rs3472489394* and *rs330736093* on *MANBA* and *SKOR2* significantly interacted with the enrichment of type IIx myonuclei in muscle and Leydig cells in the testis, respectively (Fig. 5d-e and Supplementary Fig. 19a).

**Fig. 5.**
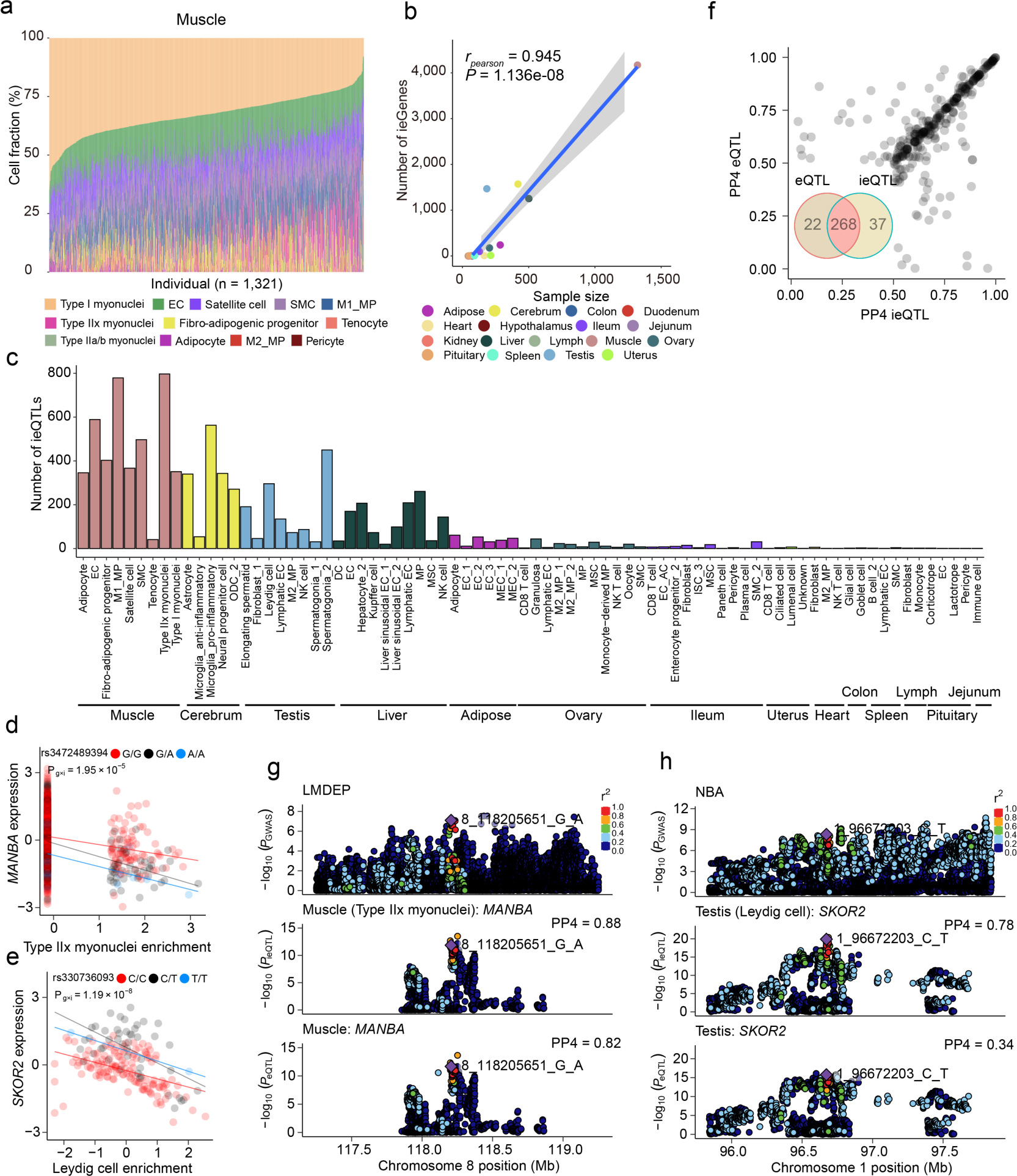
Cell-type-dependent activities of genetic variants on gene expression and pig traits. **a,** Stacked bar plots showing the fraction of cell types estimated in PigGTEx RNA-seq samples based on our snRNA-seq reference matrix in muscle tissue. **b,** Scatter plot showing the estimated number of ieGenes versus sample sizes for 17 tissues estimated using public bulk RNA-seq datasets. **c,** Number of cell type interaction QTL (ieQTL) discovered in each cell type-tissue combination at FDR < 5%. **d,** An ieQTL of *MANBA* showing cell-type-specific effects in type IIx myonuclei from muscle. Each point represents an individual and is colored by three genotypes. Both gene expression levels and cell type enrichment values are inverse normal transformed across samples. The lines are fitted by a linear regression model using the geom_smooth function from ggplot2 (v3.3.2) in R (v4.0.2). **e,** An ieQTL of *SKOR2* showing cell-type-specific effects in type IIx myonuclei from muscle. Each point represents an individual and is colored by three genotypes. Both gene expression levels and cell type enrichment values are inverse normal transformed across samples. The lines are fitted by a linear regression model using the geom_smooth function from ggplot2 (v3.3.2) in R (v4.0.2). **f,** Overlaps between ieQTL and eQTL detected by traditional bulk RNA-seq. **g,** Aligned Manhattan plots of pig GWAS, ieQTL, and eQTL at the *MANBA* locus for loin muscle depth trait (LMDEP). SNPs are colored according to the magnitude of linkage disequilibrium (*r*^2^) between adjacent SNPs pairs. **h,** Aligned Manhattan plots of pig GWAS, ieQTL, and eQTL at the *SKOR2* locus for number born alive trait (NBA). SNPs are colored according to the magnitude of linkage disequilibrium (*r*^2^) between adjacent SNPs pairs.

Furthermore, to explore the cellular effects of trait-associated variants, we performed a colocalization analysis between ieQTL and GWAS hits of 268 complex traits in pigs (Supplementary Table 2). Of the putative ieQTL, 305 loci colocalized with at least one pig GWAS hit (Fig. 5f), indicating a potential involvement in the genetic control of complex traits. By comparing GWAS colocalization results between standard PigGTEx eQTL and the newly detected ieQTL, we found that a substantial proportion of GWAS signals (> 81.96%) could be colocalized by both ieQTL and eQTL (Fig. 5f-h, Supplementary Fig. 19b, and Supplementary Table 3). For example, we found a promising colocalization between the *MANBA* gene in muscle and loin muscle depth (Fig. 5g), which was supported by both ieQTL (posterior probability of colocalization (PP4) = 0.88) and standard eQTL (PP4 = 0.82). Of note, there were 37 ieQTL-specific GWAS colocalizations (Supplementary Table 3), representing 19 complex traits, which indicated the cell-specific regulation of these traits and their potential cellular origin. We also discovered that some GWAS hits missed by bulk eQTL could be retrieved by ieQTL. A noteworthy example was the Leydig cell ieQTL of *SKOR2* in testis (Fig. 5h), which colocalized with the GWAS signal for the number of born alive at birth (PP4 = 0.78), whereas the standard eQTL from bulk testis tissues did not (PP4 = 0.34). These results together showcased the substantial potential of our cell atlas in dissecting the genetic control of the transcriptome and complex phenotypes at single-cell resolution in pigs.

### Association of cell types with complex traits and diseases in pigs and humans

Although ieQTL have provided new potential target genes and variants potentially underlying GWAS loci, the causal cell types of complex phenotypes are yet to be fully understood. To systematically infer the relevance of cell types with complex traits and diseases, we conducted the GWAS signal enrichment analyses using the signature genes of each cell type. The complex traits collected in the PigGTEx project (Supplementary Table 2) were grouped into five main categories, including reproduction traits (n = 71), health traits (n = 61), meat and carcass traits (n = 50), production traits (n = 19), and exterior traits (n = 6). Of the 263 high-resolution cell clusters in all 19 tissues, 222 (84.41%) showed significant enrichments for at least one phenotype category after multiple testing correction (Supplementary Fig. 20). For instance, the litter size relevant traits were maximally enriched in the immune cell cluster, implying the existence of critical relationships between immune function and piglet survival (Supplementary Fig. 21). Notably, many reproduction traits, such as total number born alive (NBA), total number of piglets born (TNB), and the number of stillborn pigs (NBS), showed a significant enrichments in neuronal cell types such as oligodendrocyte in cerebrum and cerebellum, in addition to Leydig cells in testis, endothelial cells in ovary and lumen cells in uterus (Fig. 6a and Supplementary Fig. 22). Moreover, several production and growth traits, including average daily gain (ADG), backfat thickness (BFT), and loin muscle area (LMA), were enriched not only in three skeletal myocytes but also in pituitary somatotropes, intestine enterocytes, and pancreatic acinar cells (Fig. 6a). However, we did not find any significant enrichment for health and exterior traits, possibly due to their relatively low GWAS power. To validate the results, we partitioned the heritability of two production traits, backfat thickness, and loin muscle depth, by cell types in a large population of over 26,000 genotyped individuals (Fig. 6b-c). As expected, we observed the enriched heritability of muscle depth trait in type IIx myonuclei. Likewise, backfat thickness showed a remarkable enrichment for enterocytes in the duodenum and enteroendocrine cells in the jejunum and colon. Although both results were obtained from two datasets with different sample sizes and distinct enrichment approaches, they showed to some extent consistency (Fig. 6a-c). Furthermore, through examining the gene-traits/disorders from Online Mendelian Inheritance in Animals database (OMIA, https://omia.org/), we identified notable cell-type-specific expression programs of many essential genes. For example, *APOE*, a major risk factor gene for Alzheimer’s disease (*47*), showed higher transcription levels in the pig astrocyte and microglia subtypes compared to other cell types. High levels of *CD163* expression (an essential receptor for the porcine reproductive and respiratory syndrome (*48*)) were mainly observed in the Kupffer cells and other macrophages.

**Fig. 6.**
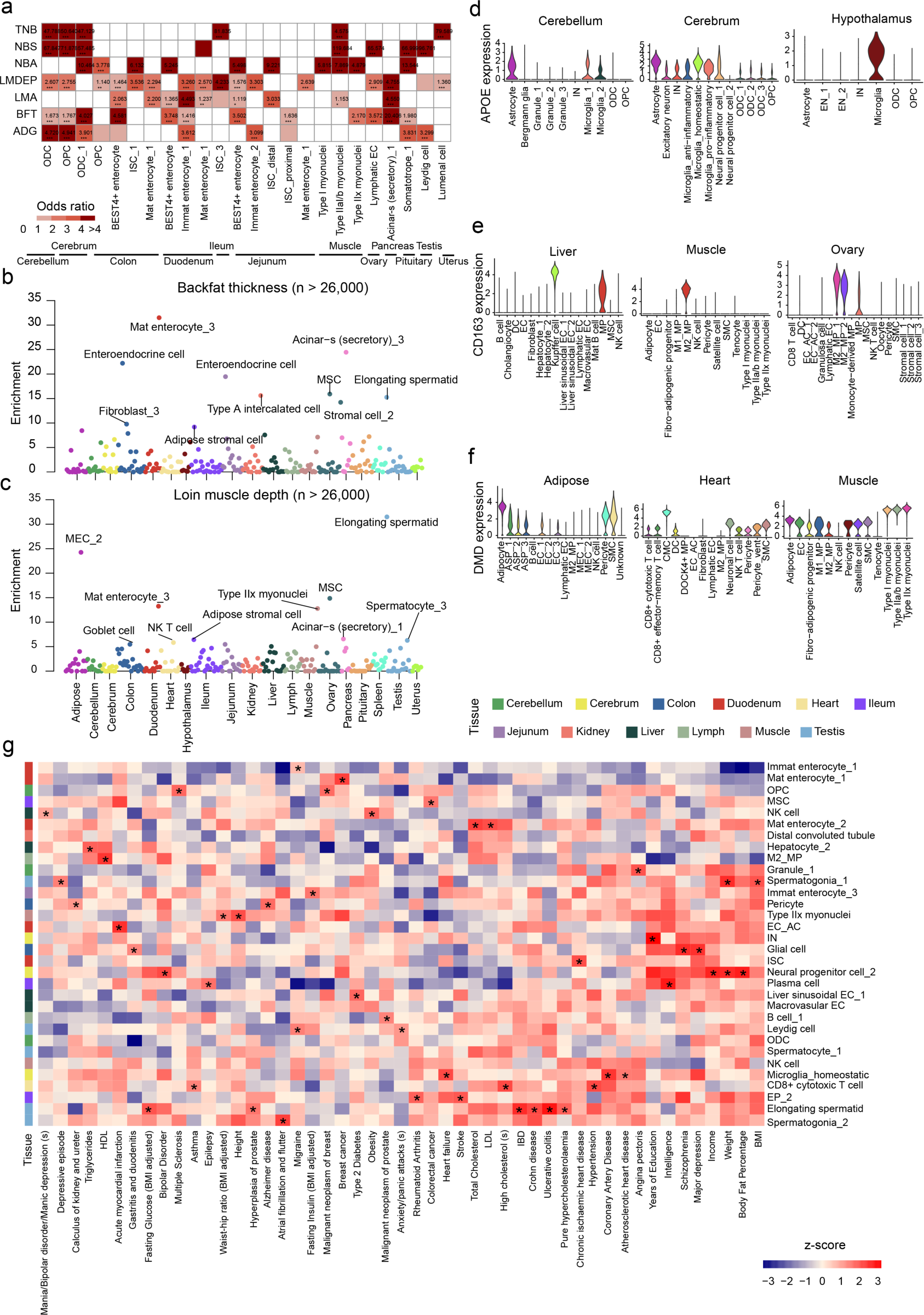
Association of single-cell transcriptomic profiles with complex traits in pigs and humans. **a,** Heatmap showing representative significant associations between cell types and traits in pigs. Definitions for abbreviations and complete results are provided in Supplementary Table 2. **b,** Evaluation of the enrichment of backfat thickness trait in putative cell types by scRNA-seq data. Each circle represents a cell-type-trait association from a large-scale population dataset. **c,** Evaluation of the enrichment of loin muscle depth trait in putative cell types by scRNA-seq data. Each circle represents a cell-type-trait association from a large-scale population dataset. **d,** Cell-type-specific expression patterns of *APOE* in the cerebellum, cerebrum, and hypothalamus. The *APOE* gene is a key candidate associated with hyperlipidemia/atherosclerosis from the OMIA database. **e,** Cell-type-specific expression patterns of *CD163* in the liver, muscle, and ovary. The *CD163* gene is an essential receptor linked to resistance/susceptibility to the porcine reproductive and respiratory syndrome (PRRS) virus from the OMIA database. **f,** Cell-type-specific expression patterns of *DMD* in the adipose, heart, and muscle. The *DMD* gene plays a vital role in muscular dystrophy from the OMIA database. **g,** Heatmap showing enrichment of pig cell types (indicated on the right) associated with selected human traits and diseases (indicated at the bottom). The colored boxes indicate selected enriched patterns. Definitions for abbreviations and complete results are provided in Supplementary Tables 4 and 5.

To explore whether our pig cell atlas could help to understand the cellular mechanisms of complex traits and diseases in humans, we quantified the heritability enrichment of 137 human complex phenotypes (Supplementary Table 4) across the 261 annotated cell types (two cell clusters defined as unknown types were discarded) via the stratified linkage disequilibrium score regression analysis (LDSC). We retrieved 15,354 one-to-one pig-human orthologous protein-coding genes from the Ensembl dataset (version 104) for the following analyses. Our results revealed a total of 1,547 significant associations (the corrected enrichment *p*-value < 0.05) between pig cell types and human complex phenotypes (Fig. 6g, Supplementary Fig. 23, and Supplementary Table 5). As expected, we observed significant enrichments of several neurological and psychiatric phenotypes, such as multiple sclerosis, schizophrenia, and bipolar disorder, in neural cell types, including excitatory neurons and neural progenitor cells from the cerebrum, as well as in certain immune cell clusters such as microglia from cerebrum and macrophages from pituitary. Additionally, metabolic traits, including type 2 diabetes and cholesterol-related phenotypes, showed expected associations with hepatocytes, pancreatic duct cells, and ileum goblet cells, as well as interesting associations with several skeletal muscle and intestine cell populations. Moreover, our analysis revealed some novel relationships between GWAS traits and cell types. For instance, we found enriched heritability of several intestine diseases, such as Crohn’s disease and diverticular disease, in cell clusters corresponding to brain-resident immune cells (*5, 15*), in addition to enterocytes and immune cells from the four intestine segments. For fasting insulin and glucose traits, we found significant enrichments in adipocytes from adipose and skeletal muscle cells and enterocytes from the intestines. Similarly, we observed striking enrichments of anthropometric traits, including height, waist-hip ratio, and body fat percentage, not only in intestinal stem cells, fibro-adipogenic progenitor cells from skeletal muscle, and adipocyte from adipose but also in multiple cell populations from testis and ovary. Overall, our pig snRNA-seq data provided new comprehensive insights into trait-relevant cell types in both pigs and humans, which will boost the unraveling of molecular and cellular mechanisms underlying complex phenotypes and the potential utilization of pigs as human biomedical models for certain diseases.

## Discussion

The domestic pig (*Sus scrofa*) is a valuable livestock species that contributes significantly to both agricultural and biomedical research. Recent studies, including our PigGTEx project, have revealed that many traits-associated variants are located in non-coding regions and affect the spatiotemporal expression of candidate genes in a context-specific (tissue- or cell-type-specific) fashion. However, the impacts of genetic variations on these regulatory pathways and how they vary across trait-relevant cell types have not been explored in pigs. To bridge the gaps between genetic variants and phenotypes at single-cell resolution, we performed a comprehensive analysis by integrating a cross-tissue snRNA-seq atlas with the large-scale PigGTEx datasets. This work not only establishes a comprehensive single-cell reference map as a baseline for dissecting cellular heterogeneity within and across tissues but also highlights a more powerful strategy for identifying trait-critical cellular signatures and cell-type-specific eQTL in pigs.

The present study employed single-nucleus RNA-seq to profile gene expression in 229,268 high-quality cells from 19 tissues in pigs, similar to a recent study (*27*) which constructed the first single-cell transcriptomic atlas of 222,526 cells across 20 swine tissues. Compared with that work, our dataset represents a broader range of pig organ sources covering nine major body systems and especially comprises several highly important tissues in pig production performance, such as skeletal muscle, four intestine segments, and three reproductive organs. Given the large diversity in the chosen tissues, the two studies demonstrate a good complement and represent very significant contributions to the efforts of the pig single-cell consortium. In line with single-cell landscapes in other species (*1, 2, 4, 5, 49-51*), we identified primary cell classes based on known canonical marker genes and captured a few rare cell types like Purkinje cells from the brain and enteroendocrine cells from the intestine, which may facilitate our understanding of cell lineage trajectory and tissue homeostasis. Our pig cross-tissue cell atlases clarify the heterogeneous characteristics in cellular compositions and molecular properties within and across tissues. For example, we delineated the global transcriptional divergence and transition pattern among three dominant myofiber types (type I, IIa/b, and IIx) and revealed evolutionarily conserved similarity in pivotal genes specializing myofiber, such as *MYH7* and *MYBPC2* across mammals (*35, 36, 52*). This finding may have important implications for improving meat quality and quantity, which are largely affected by myofiber characteristics and proportions in pigs (*53, 54*). Type II myonuclei exhibited a notable enrichment in metabolic processes, indicating their crucial involvement in metabolic traits and syndromes, *i.e.*, meat production and fat deposition in pigs and type 2 diabetes and obesity in humans (*35, 55–57*). Our data also demonstrate the prevalence of epithelial and immune cells across different tissue contexts and offer a more detailed understanding of cell compartments. Although some cells of a common type are shared across tissues, subpopulations are specifically enriched in particular tissues. These tissue-resident epithelial and immune subsets are specialized to fulfill the specific functional demands of different tissues, probably owing to unique local environments or niches (*50, 58*).

Although our PigGTEx project has provided a compendium of genetic regulatory effects across pig tissues and functional variants underlying complex traits (*24*), a comprehensive understanding of gene regulation at the single-cell resolution for most non-coding loci is still lacking. To address this issue, emerging approaches such as single-cell eQTL and heritability enrichment analyses have been extensively used in deciphering complex human traits and diseases (*14–17, 46*) but have yet to be systematically applied in pig studies. As a critical component of the PigGTEx project, our work offers an in-depth dissection of the genetic effects of trait-critical cellular signatures and cell-type-specific eQTL, in addition to the comprehensive pig cell reference map, setting it apart from other recent single-cell studies (*27*). We revealed that around 15% of the loci that co-localized with GWAS traits showed significant cell-type specificity, underscoring the advantages of single-cell eQTL analysis over the standard bulk eQTL approach. The proportion missed by bulk studies is slightly lower than what has been described in humans (*46*), which might be attributed to the limited sample size in our work. By linking individual cell types to complex traits, we identified substantial cell-type-trait associations that are consistent with previous studies (*5, 15, 16, 35*), suggesting high functional conservation of major cell types among mammal species (*52*). Furthermore, we mapped several unique associations between cell types and important phenotypes in pigs, such as the driving roles of myofiber cell types for meat production traits and Leydig cells from the testis for reproduction traits. Overall, our results provide meaningful insights into previously cryptic molecular and cellular mechanisms behind traits of economic importance and offer new opportunities for precision breeding in pigs.

Despite the significant findings of our study, several limitations must be noted. Firstly, the current dataset comprises only one male and one female pig and is not an exhaustive characterization of all pig organs. As such, we cannot fully capture the complete single-cell picture and inter-individual variation in cellular composition, potentially limiting our ability to explore rare cell types and map entire trait-associated cellular signatures. Secondly, compared with single-cell RNA-seq, our single-nucleus RNA-seq can only profile nuclear transcripts and not cytoplasmic transcripts. Different library preparation protocols may result in a reasonable proportion difference in specific cell types, such as muscle, neural, and immune cells, despite globally consistent detection performance in gene number and cell type diversity between them (*59, 60*). Lastly, the sample size of certain tissues used in cellular deconvolution and heritability partitioning analyses is relatively small, limiting the statistical power to detect causative trait-associated cell types and single-cell eQTL. Therefore, future studies that incorporate larger sample sizes, a broader range of tissues, and multiple complementary single-cell approaches will be required to provide robust evidence and facilitate a more comprehensive interpretation of our findings.

In summary, this study presents a compendium of high-resolution body-wide single-cell transcriptional landscape, provides a deeper understanding of the expression patterns and functions of tissue-specific and shared cell types, and illuminates the intricate cell-cell interactions governing tissue homeostasis. Through pioneering single-cell eQTL and colocalization analyses in pigs, we pinpointed the likely causative cell-type-associated variants and genes underlying traits of economic importance. Additionally, thousands of cell-type-trait associations were discovered, and previously unexplored biological mechanisms were explicated using heritability enrichment analysis. Collectively, these findings will significantly enhance our understanding of cross-tissue and cross-individual variations of cellular phenotypes and highlight promising trait-associated determinants (variants and cell types) for advancing the fields of future pig breeding and human biomedical research.

## Methods

### Ethics statement

All animal protocols and procedures were implemented in compliance with the Guide for the Care and Use of Experimental Animals established by the Ministry of Agriculture and Rural Affairs (Beijing, China) and were approved by the Institutional Animal Care and Use Committee of the Chinese Academy of Agricultural Sciences. Prior to tissue sampling, the pigs were humanely euthanized as necessary to minimize their suffering.

### Tissue collection and single-nucleus suspension

One male and one female Meishan pig, aged 180 days, were obtained from a commercial pig farming company managed under the same conditions (Nantong, Jiangsu). Nineteen tissues, including adipose, cerebellum, cerebrum, colon, duodenum, heart, hypothalamus, ileum, jejunum, kidney, liver, lymph, muscle, ovary, pancreas, pituitary, spleen, testis, and uterus, were freshly harvested from postmortem samples. Each tissue was kept on ice and minced into 5-10 pieces weighing approximately 50-100 mg each on ice with sterilized scissors. Tissue samples were then snap-frozen in liquid nitrogen and stored at −80℃ until nuclear extraction was performed. Single-nucleus isolation was conducted as previously described (*28, 59*). Briefly, tissue samples were homogenized using the Dounce homogenizer with 25 strokes of the loose pestle A and then 25 strokes of the tight pestle B in 1 ml of ice-cold homogenization buffer. After this, the mixture was filtered through a 40-μm cell strainer into a 1.5-ml tube. To collect dissociated single nuclei, the sample was centrifuged at 500g for 5 min at 4℃, and the supernatant was discarded. After centrifugation, the nuclear pellet was resuspended using an appropriate amount of 1× PBS/0.5% BSA with RNase inhibitor, filtered through a 40-μm cell strainer, and counted. A final concentration of 1,000 nuclei per µl was used for library preparation.

### Single-nucleus RNA-seq library preparation and sequencing

The single-nucleus RNA-seq libraries were prepared following the standard protocol supplied by 10× Genomics (Berry Genomics, Beijing, China). In brief, isolated nuclei were captured in droplets with gel beads in the Chromium Controller. Following the RNA reverse transcription step, emulsions were broken, and barcoded cDNA was purified with Dynabeads, after which PCR amplification was performed. The amplified cDNA was then used for 3’ gene expression library construction. Then, indexed libraries were constructed according to the manufacturer’s recommendations. After quality control, eligible libraries were sequenced on the Novaseq 6000 platform (Illumina) in a 150 bp paired-end manner. The first 28 bp in read 1 captured both the 16 bp 10× barcode and the 12 bp UMI.

### Preprocessing of snRNA-seq data

The Sscrofa11.1 reference assembly (*61*) in FASTA format and annotated gene model in GTF format were downloaded from the Ensembl database (ftp://ftp.ensembl.org/pub/release-101/). Raw snRNA-seq data were aligned to the pig reference genome and subjected to barcode assignment and unique molecular identifier (UMI) counting using the commands recommended by the CellRanger pipeline (10× Genomics, CA, USA). Given that snRNA-seq captures both unspliced pre-mRNA and mature mRNA, we used the *include-introns* option for counting exonic and intronic reads together. The filtered gene expression matrix was used for further analysis with the Seurat package (*62*). To ensure the accuracy and robustness of our results, we removed ambient RNA and potential doublets using DecontX (*63*) and DoubletFinder (*64*) with default settings. We also filtered out low-quality nuclei expressing less than 200 genes or more than 5,000 genes, and less than 500 UMIs or more than 15,000 UMIs, as well as those exceeding 5% of mitochondrial content. During the gene filter step, all genes not expressed in at least three nuclei were removed. In addition, to balance our dataset in subsequent analyses, we randomly selected 20,000 nuclei from the spleen, as it had a much higher number (n = 53,444) of captured nuclei compared to other tissues.

### Cell clustering and cell type annotation

After filtering, the remaining high-quality data were log-normalized and scaled to account for cell-to-cell variation with regression on the number of UMIs and percentage of mitochondrial genes. Subsequently, PCA linear dimensionality reduction analysis was performed, followed by t-Distributed Stochastic Neighbor Embedding (t-SNE) and Uniform Manifold Approximation and Projection (UMAP) visualization approaches using the Scanorama tool (*65*), to capture the global cell type landscapes across tissues. For individual clustering, each tissue dataset was visualized using the Seurat package (*62*). Parameters used in each function were manually curated to obtain the optimal clustering of cells by adjusting the number of principal components and resolutions on a per-dataset basis. We employed the *FindAllMarkers* or *FindMarkers* function with default parameters to identify marker genes of each cluster and annotated each cell type based on known classical markers from extensive published literature. The Pearson or Spearman correlation coefficients among cell types were calculated using the average expression of the top 1,000 highly variable features, and we used the pheatmap package (https://github.com/raivokolde/pheatmap) to visualize the results. Besides, the expression of marker genes in different cell types was visualized with the ggplot2 R package.

### Cell type diversity estimation

Shannon entropy was calculated to evaluate cell type diversity in each tissue with a previously published method (*14*) according to the formula − ∑*_x_*(*p_x_* × *log*_2_(*p_x_*)), where *p_x_* is the proportion of each cell type *x* in a tissue. The entropy value per tissue was plotted using the ggplot2 R package.

### Pseudotime trajectory inference and RNA velocity analysis

The cell lineage trajectory was inferred using Monocle 3 (*41*) according to the standard tutorial. We used the built-in DDRTree algorithm for dimensional reduction and visualization after constructing the cell trajectory. Notably, the root state of the inferred trajectory was specified based on existing biological knowledge. Furthermore, we predicted the velocity streams and latent time assignments from sorted bam files using the dynamical model implemented in scVelo (*66*).

### Cell cycle index estimation

To further infer dynamic information about cell state, we calculated a cell cycle index for each cell type with a previously published method (*3*). Typically, progenitor cells with rapidly dividing capacity display higher cycling indices, whereas cell types that are known to be largely quiescent exhibit lower values.

### Cell-cell interaction analysis

To investigate cellular communication patterns between different cell types, we used the CellChat package (*42*) with default parameters, which is a manually curated database of literature-supported ligand-receptor interactions in humans and mice. To run CellChat analysis in pig datasets, we mapped pig gene symbols to human orthologs. Ligand-receptor pairs with *p*-value < 0.05 were considered to be significant.

### Tissue enrichment of clusters

We estimated the enrichment of each cell cluster across tissues, as previously described (*67*). In brief, we calculated the observed and expected cell numbers in each cell cluster to compute the ratio (R_o/e_) between the two values using the epitools R package. We considered a cluster to be enriched in a specific tissue if R_o/e_ > 1.

### Gene ontology (GO) enrichment analysis

Gene Ontology (GO) analysis was performed using the clusterProfiler 4.0 (*68*) and org.Hs.eg.db annotation package, considering that genome-wide annotation is incomplete in pigs. The Benjamini-Hochberg (BH) procedure was used for the multiple testing corrections, and only GO terms with an adjusted *p*-value smaller than 0.05 were retained.

### Single-cell regulatory network analysis

To uncover cell-type-specific transcription regulons and construct gene regulation networks (GRNs), we conducted single-cell regulatory network inference and clustering analysis using the SCENIC suite (*69*) with the default parameters. The original expression matrix was normalized with Seurat and fed into SCENIC to build a coexpression network using the built-in GRNBoost2 algorithm. The activity of regulons in each cell was calculated by the AUCell algorithm.

### Cellular deconvolution analysis using CIBERSORT

For each tissue, we first identified differentially expressed genes specific to each cell type using the *Findmarkers* function in the Seurat package. We then selected the top 50 genes with the most significant overexpression, based on adjusted *p*-value (< 0.05) and average log_2_ fold change (> 0.5), to build the gene expression signature matrix for the cell-type reference set. To predict the abundances of cell types in a mixed cell population for each tissue, we collected the RNA-seq expression matrix of 17 matching bulk tissues with our snRNA-seq data from the PigGTEx database (http://piggtex.farmgtex.org/). Subsequently, the CIBERSORT tool (*44*) was selected for cellular deconvolution analysis, given its great resolving power (*70*). To test the robustness of cellular deconvolution, we first used the *generateBulk_norep* function in the SCDC package (*45*) to obtain the transcript per million (TPM) matrix of 1,000 pseudo bulk samples (default parameters) with known cell type distribution based on our liver snRNA-seq data. Then we used CIBERSORT to perform deconvolution on these samples using the TPM matrix of signature genes from each cell type in pig liver. The number of permutation tests was set to 1,000 times to determine the significance level, and p < 0.05 was regarded as statistical significance. Finally, we calculated the Spearman correlation coefficient between the known and predicted cell type distribution of hepatocyte cells to assess the accuracy of CIBERSORT deconvolution in our pig dataset.

### Cell type interaction *cis*-eQTL mapping

To detect whether a *cis*-eQTL explicitly affects gene expression in a given cell type, we performed cell type interaction *cis*-eQTL (ieQTL) mapping for 17 bulk tissues of PigGTEx. We used the cell type composition (i.e., enrichment score) estimated from CIBERSORTx as above and only considered cell types with an enrichment score > 0 in at least 20 samples and/or 20% of samples within a tissue. For each tissue-cell type pair, we performed ieQTL mapping via a linear regression model implemented in TensorQTL (v1.0.3) (*71*), which included an interaction term between genotype and cell type enrichment score:

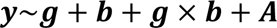

where ***y*** is the vector of gene expression values (*i.e.*, the inverse normal transformed TMM), ***g*** is the genotype dosage (*i.e.*, 0/1/2) vector of the tested SNP from PigGTEx samples, ***b*** is the enrichment score of a given cell type predicted from snRNA-seq data, ***g*** × ***b*** is the interaction term between genotype and enrichment score, and A represents the covariates (*i.e.*, genotype PCs and PEER factors, detailed in PigGTEx pilot phase). For the ieQTL mapping, we only considered SNPs within ±1 Mb of transcription start sites (TSS) of each gene. We eliminated those SNPs with minor allele frequency (MAF) < 0.05 in the top and/or bottom 50% of samples sorted by the enrichment score of each cell type, using TensorQTL (v1.0.3) with parameter: -- maf_threshold_interaction 0.05. To correct for the multiple testing at the gene level, we used eigenMT (*72*) in TensorQTL for calculating the top nominal p-value of each gene. We then computed the genome-wide significance of genes using the Benjamini-Hochberg FDR correction on the eigenMT-corrected *p*-values and defined as ieGene that with at least one significant ieQTL (*i.e.*, FDR-corrected *p*-value < 0.05).

### Allele-specific expression validation of ieQTL

We used allele-specific expression (ASE) data at the individual level to validate the discovered ieQTL. We first estimated the effect size (i.e., allelic fold change, aFC) of the top ieQTL for each ieGene from ASE data using the script phaser_cis_var.py in phASER (v1.1.1) (*73*) and considered only ieQTL with nominally significant ASE (*p*-value < 0.05) data in more than ten heterozygous individuals with more than eight reads for a gene. To filter out outlier samples in ASE data, we applied the median absolute deviation (MAD) based on Hampel’s test to the allelic imbalance (AI) ratio values 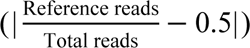 across samples (*46, 74*). When a sample had |AI_i_ − median (AI)| ≥ 4.5 × MAD, where MAD = median(|AI_i_ − median(AI)|) and AI_#_ is the allelic imbalance ratio value for the ith individual, we defined it as an outlier and eliminated it in the validation process. Within a given tissue, we determined that an ieQTL was validated by ASE data if it presented a nominally significant (*p*-value < 0.05) Pearson’s correlation between allelic fold change (aFC) of an ASE locus and cell type enrichment score across samples.

### Colocalization between ieQTL and GWAS loci

To identify shared association variants between the ieQTL and GWAS loci retrieved from the PigGTEx project, we performed colocalization analysis using the Bayesian statistical procedure implemented in Coloc (v5.1) (*75*). Briefly, we used the summary statistics of ieQTL for each ieGene and its matched GWAS loci as input for Coloc. We only considered the GWAS loci with at least one SNP with a *p*-value < 1×10^-5^. We obtained posterior probabilities PP4 from the *coloc.abf* function with default parameters, where PP4 represents the probabilities of a shared variant affecting both the gene expression of a given cell type and the complex trait. We defined ieGene-trait pairs with PP4 > 0.5 as significant colocalization. In addition, to compare whether eQTL differ from ieQTL in terms of colocalization with GWAS loci, we used the same pipeline employed for ieQTL scanning to perform the colocalization analysis for eQTL for each ieGene and its matched GWAS loci as well.

### Genetic mapping of cell type specificity for complex traits in pigs and humans

To uncover associations of traits with cell types, we performed an enrichment analysis of significant GWAS loci and cell-type-specific genes using the LOLA (v1.22.0) R package (*76*). Specifically, we extracted the top 200 cell-type-specific genes sorted in ascending order by the *p*-value for each of the 19 available tissues and created an annotation based on the genomic regions of these candidate genes for each cell type-tissue pair. We then examined the GWAS summary statistics of 268 pig complex traits and selected significant genetic variants with *p* < 5 × 10^-8^ for each trait (*24*), using all tested SNPs of the 268 GWAS summaries as the background set. Finally, we calculated the significance level (*p*-value) of the enrichment fold using Fisher’s exact test with FDR correction and defined trait-tissue-cell type trios with *p*-value < 0.05 as significant enrichment. Furthermore, we expanded our enrichment analysis to a larger Duroc population (> 26,000 individuals) from a commercial company, given that the current GWAS dataset is relatively small. We performed heritability enrichment analysis for backfat thickness and loin muscle depth traits with genomic partitioning of quantitative genetic variance similar to (*77*). A total of 11.7 M imputed variants that had been quality-controlled were grouped into two sets: one containing variants within ±10 Kb of the top genes specific to each cell type, and the other containing the remaining variants. Per-variant heritability enrichment was calculated for each cell type-specific variant set.

To test the enrichment of genes associated with human traits and diseases for each specific pig cell type, we collected the GWAS summary statistics of 137 human complex traits from the UK Biobank and public literature (Supplementary Table 4). We converted cell-type-specific genes in pigs to the corresponding human orthologous genes with one-to-one mapping with the Ensembl database. Finally, we employed linkage disequilibrium (LD) score regression analysis (https://github.com/bulik/ldsc) (*78, 79*) to partition the heritability based on 262 annotations, including 261 cell-type-specific gene lists and one base annotation including all SNPs. Heritability enrichment was calculated as the proportion of trait heritability contributed by SNPs in the annotation over the total proportion of SNPs in that annotation. We reported the coefficient *p*-value as a measure of the association of each cell type with the traits. All plots showed the predicted z-score of partitioned LD score regression.

### Statistics and reproducibility

If not specified, all statistical analyses and data visualization were performed in the R environment. No statistical method was used to predetermine sample size, no data were excluded from the analyses, and all analyses were not randomized, ensuring maximum reproducibility.

## Data availability

Raw sequencing reads generated by this work were deposited in the National Center for Biotechnology Information database under the accession number GSE233285. Analysis codes in this work are available at https://github.com/chenlijuan009/PigCellAtlas.

## Acknowledgments

We thank Prof. Elisabetta Giuffra (INRAE, France) for her valuable comments and suggestions on this project. This work was supported by the National Key Research and Development Program of China (2021YFF1000600), the National Natural Science Foundation of China (32002150 to G.Y., 32022078 to Z.Z.), the Basic and Applied Basic Research Foundation of Guangdong Province (2020B1515120053 to G.Y.), the Shenzhen Science and Technology Innovation Commission (JCYJ20190813114401691 to G.Y.), the Local Innovative and Research Teams Project of Guangdong Province (2019BT02N630 to Z.Z.), the China Agriculture Research System (CARS-35 to Z.Z. and Jiaqi L.), the Key Research and Development projects in Yangzhou City (YZ2021037 to Z.C.), the Independent Research Fund Denmark (8048-00072A to L.L.), the Novo Nordisk Foundation (NNF21OC0071718 to L.L., NNF21OC0071031 and NNF21OC0068988 to Y.L.), and the Lundbeck Foundation (R396-2022-350 to Y.L.).

## Author contributions

G.Y. and L.F. conceived and designed the study. X.Q., Z.C., and L.Y. were responsible for sample collection. L.C., H.L., J.T., Z.W., X.C., Jinghui L. H.Z., Z.B., and J.J. conducted bioinformatic analysis. L.C. and Z.W. performed snRNA-seq analyses. H.L., X.P., and J.T. contributed to eQTL mapping and cellular deconvolution. J.T., X.C., Jinghui L., J.J., Z.Z., and Jiaqi L. were responsible for GWAS data collection and analysis in pigs and humans. G.Y., H.L., J.T., and Jinghui L. wrote the initial draft of the manuscript. G.Y., L.F., J.J., G.E.L., F.W., L.L., Y.L., G.S., M.S.L., M.B., D.C.P., P.K.M., M.F., A.C., M.A., C.L., C.K.T., and O.M. revised the manuscript. All authors read and approved the final manuscript.

## References

1. X. Han, R. Wang, Y. Zhou, L. Fei, H. Sun, S. Lai, A. Saadatpour, Z. Zhou, H. Chen, F. Ye, D. Huang, Y. Xu, W. Huang, M. Jiang, X. Jiang, J. Mao, Y. Chen, C. Lu, J. Xie, Q. Fang, Y. Wang, R. Yue, T. Li, H. Huang, S. H. Orkin, G. C. Yuan, M. Chen, G. Guo, Mapping the Mouse Cell Atlas by Microwell-Seq. Cell 173, 1307 (2018).

2. X. Han, Z. Zhou, L. Fei, H. Sun, R. Wang, Y. Chen, H. Chen, J. Wang, H. Tang, W. Ge, Y. Zhou, F. Ye, M. Jiang, J. Wu, Y. Xiao, X. Jia, T. Zhang, X. Ma, Q. Zhang, X. Bai, S. Lai, C. Yu, L. Zhu, R. Lin, Y. Gao, M. Wang, Y. Wu, J. Zhang, R. Zhan, S. Zhu, H. Hu, C. Wang, M. Chen, H. Huang, T. Liang, J. Chen, W. Wang, D. Zhang, G. Guo, Construction of a human cell landscape at single-cell level. Nature 581, 303–309 (2020).

3. C. Tabula Sapiens, R. C. Jones, J. Karkanias, M. A. Krasnow, A. O. Pisco, S. R. Quake, J. Salzman, N. Yosef, B. Bulthaup, P. Brown, W. Harper, M. Hemenez, R. Ponnusamy, A. Salehi, B. A. Sanagavarapu, E. Spallino, K. A. Aaron, W. Concepcion, J. M. Gardner, B. Kelly, N. Neidlinger, Z. Wang, S. Crasta, S. Kolluru, M. Morri, A. O. Pisco, S. Y. Tan, K. J. Travaglini, C. Xu, M. Alcantara-Hernandez, N. Almanzar, J. Antony, B. Beyersdorf, D. Burhan, K. Calcuttawala, M. M. Carter, C. K. F. Chan, C. A. Chang, S. Chang, A. Colville, S. Crasta, R. N. Culver, I. Cvijovic, G. D’Amato, C. Ezran, F. X. Galdos, A. Gillich, W. R. Goodyer, Y. Hang, A. Hayashi, S. Houshdaran, X. Huang, J. C. Irwin, S. Jang, J. V. Juanico, A. M. Kershner, S. Kim, B. Kiss, S. Kolluru, W. Kong, M. E. Kumar, A. H. Kuo, R. Leylek, B. Li, G. B. Loeb, W. J. Lu, S. Mantri, M. Markovic, P. L. McAlpine, A. de Morree, M. Morri, K. Mrouj, S. Mukherjee, T. Muser, P. Neuhofer, T. D. Nguyen, K. Perez, R. Phansalkar, A. O. Pisco, N. Puluca, Z. Qi, P. Rao, H. Raquer-McKay, N. Schaum, B. Scott, B. Seddighzadeh, J. Segal, S. Sen, S. Sikandar, S. P. Spencer, L. C. Steffes, V. R. Subramaniam, A. Swarup, M. Swift, K. J. Travaglini, W. Van Treuren, E. Trimm, S. Veizades, S. Vijayakumar, K. C. Vo, S. K. Vorperian, W. Wang, H. N. W. Weinstein, J. Winkler, T. T. H. Wu, J. Xie, A. R. Yung, Y. Zhang, A. M. Detweiler, H. Mekonen, N. F. Neff, R. V. Sit, M. Tan, J. Yan, G. R. Bean, V. Charu, E. Forgo, B. A. Martin, M. G. Ozawa, O. Silva, S. Y. Tan, A. Toland, V. N. P. Vemuri, S. Afik, K. Awayan, O. B. Botvinnik, A. Byrne, M. Chen, R. Dehghannasiri, A. M. Detweiler, A. Gayoso, A. A. Granados, Q. Li, G. Mahmoudabadi, A. McGeever, A. de Morree, J. E. Olivieri, M. Park, A. O. Pisco, N. Ravikumar, J. Salzman, G. Stanley, M. Swift, M. Tan, W. Tan, A. J. Tarashansky, R. Vanheusden, S. K. Vorperian, P. Wang, S. Wang, G. Xing, C. Xu, N. Yosef, M. Alcantara-Hernandez, J. Antony, C. K. F. Chan, C. A. Chang, A. Colville, S. Crasta, R. Culver, L. Dethlefsen, C. Ezran, A. Gillich, Y. Hang, P. Y. Ho, J. C. Irwin, S. Jang, A. M. Kershner, W. Kong, M. E. Kumar, A. H. Kuo, R. Leylek, S. Liu, G. B. Loeb, W. J. Lu, J. S. Maltzman, R. J. Metzger, A. de Morree, P. Neuhofer, K. Perez, R. Phansalkar, Z. Qi, P. Rao, H. Raquer-McKay, K. Sasagawa, B. Scott, R. Sinha, H. Song, S. P. Spencer, A. Swarup, M. Swift, K. J. Travaglini, E. Trimm, S. Veizades, S. Vijayakumar, B. Wang, W. Wang, J. Winkler, J. Xie, A. R. Yung, S. E. Artandi, P. A. Beachy, M. F. Clarke, L. C. Giudice, F. W. Huang, K. C. Huang, J. Idoyaga, S. K. Kim, M. Krasnow, C. S. Kuo, P. Nguyen, S. R. Quake, T. A. Rando, K. Red-Horse, J. Reiter, D. A. Relman, J. L. Sonnenburg, B. Wang, A. Wu, S. M. Wu, T. Wyss-Coray, The Tabula Sapiens: A multiple-organ, single-cell transcriptomic atlas of humans. Science 376, eabl4896 (2022).

4. Y. Liao, L. Ma, Q. Guo, W. E, X. Fang, L. Yang, F. Ruan, J. Wang, P. Zhang, Z. Sun, H. Chen, Z. Lin, X. Wang, X. Wang, H. Sun, X. Fang, Y. Zhou, M. Chen, W. Shen, G. Guo, X. Han, Cell landscape of larval and adult Xenopus laevis at single-cell resolution. Nature Communications 13, 4306 (2022).

5. L. Han, X. Wei, C. Liu, G. Volpe, Z. Zhuang, X. Zou, Z. Wang, T. Pan, Y. Yuan, X. Zhang, P. Fan, P. Guo, Y. Lai, Y. Lei, X. Liu, F. Yu, S. Shangguan, G. Lai, Q. Deng, Y. Liu, L. Wu, Q. Shi, H. Yu, Y. Huang, M. Cheng, J. Xu, Y. Liu, M. Wang, C. Wang, Y. Zhang, D. Xie, Y. Yang, Y. Yu, H. Zheng, Y. Wei, F. Huang, J. Lei, W. Huang, Z. Zhu, H. Lu, B. Wang, X. Wei, F. Chen, T. Yang, W. Du, J. Chen, S. Xu, J. An, C. Ward, Z. Wang, Z. Pei, C. W. Wong, X. Liu, H. Zhang, M. Liu, B. Qin, A. Schambach, J. Isern, L. Feng, Y. Liu, X. Guo, Z. Liu, Q. Sun, P. H. Maxwell, N. Barker, P. Munoz-Canoves, Y. Gu, J. Mulder, M. Uhlen, T. Tan, S. Liu, H. Yang, J. Wang, Y. Hou, X. Xu, M. A. Esteban, L. Liu, Cell transcriptomic atlas of the non-human primate Macaca fascicularis. Nature 604, 723–731 (2022).

6. J. Lonsdale, J. Thomas, M. Salvatore, R. Phillips, E. Lo, S. Shad, R. Hasz, G. Walters, F. Garcia, N. Young, B. Foster, M. Moser, E. Karasik, B. Gillard, K. Ramsey, S. Sullivan, J. Bridge, H. Magazine, J. Syron, J. Fleming, L. Siminoff, H. Traino, M. Mosavel, L. Barker, S. Jewell, D. Rohrer, D. Maxim, D. Filkins, P. Harbach, E. Cortadillo, B. Berghuis, L. Turner, E. Hudson, K. Feenstra, L. Sobin, J. Robb, P. Branton, G. Korzeniewski, C. Shive, D. Tabor, L. Qi, K. Groch, S. Nampally, S. Buia, A. Zimmerman, A. Smith, R. Burges, K. Robinson, K. Valentino, D. Bradbury, M. Cosentino, N. Diaz-Mayoral, M. Kennedy, T. Engel, P. Williams, K. Erickson, K. Ardlie, W. Winckler, G. Getz, D. DeLuca, D. MacArthur, M. Kellis, A. Thomson, T. Young, E. Gelfand, M. Donovan, Y. Meng, G. Grant, D. Mash, Y. Marcus, M. Basile, J. Liu, J. Zhu, Z. Tu, N. J. Cox, D. L. Nicolae, E. R. Gamazon, H. K. Im, A. Konkashbaev, J. Pritchard, M. Stevens, T. Flutre, X. Wen, E. T. Dermitzakis, T. Lappalainen, R. Guigo, J. Monlong, M. Sammeth, D. Koller, A. Battle, S. Mostafavi, M. McCarthy, M. Rivas, J. Maller, I. Rusyn, A. Nobel, F. Wright, A. Shabalin, M. Feolo, N. Sharopova, A. Sturcke, J. Paschal, J. M. Anderson, E. L. Wilder, L. K. Derr, E. D. Green, J. P. Struewing, G. Temple, S. Volpi, J. T. Boyer, E. J. Thomson, M. S. Guyer, C. Ng, A. Abdallah, D. Colantuoni, T. R. Insel, S. E. Koester, A. R. Little, P. K. Bender, T. Lehner, Y. Yao, C. C. Compton, J. B. Vaught, S. Sawyer, N. C. Lockhart, J. Demchok, H. F. Moore, The Genotype-Tissue Expression (GTEx) project. Nat. Genet. 45, 580–585 (2013).

7. F. Aguet, S. Anand, K. G. Ardlie, S. Gabriel, G. A. Getz, A. Graubert, K. Hadley, R. E. Handsaker, K. H. Huang, S. Kashin, X. Li, D. G. MacArthur, S. R. Meier, J. L. Nedzel, D. T. Nguyen, A. V. Segrè, E. Todres, B. Balliu, A. N. Barbeira, A. Battle, R. Bonazzola, A. Brown, C. D. Brown, S. E. Castel, D. F. Conrad, D. J. Cotter, N. Cox, S. Das, O. M. de Goede, E. T. Dermitzakis, J. Einson, B. E. Engelhardt, E. Eskin, T. Y. Eulalio, N. M. Ferraro, E. D. Flynn, L. Fresard, E. R. Gamazon, D. Garrido-Martín, N. R. Gay, M. J. Gloudemans, R. Guigó, A. R. Hame, Y. He, P. J. Hoffman, F. Hormozdiari, L. Hou, H. K. Im, B. Jo, S. Kasela, M. Kellis, S. Kim-Hellmuth, A. Kwong, T. Lappalainen, X. Li, Y. Liang, S. Mangul, P. Mohammadi, S. B. Montgomery, M. Muñoz-Aguirre, D. C. Nachun, A. B. Nobel, M. Oliva, Y. Park, Y. Park, P. Parsana, A. S. Rao, F. Reverter, J. M. Rouhana, C. Sabatti, A. Saha, M. Stephens, B. E. Stranger, B. J. Strober, N. A. Teran, A. Viñuela, G. Wang, X. Wen, F. Wright, V. Wucher, Y. Zou, P. G. Ferreira, G. Li, M. Melé, E. Yeger-Lotem, M. E. Barcus, D. Bradbury, T. Krubit, J. A. McLean, L. Qi, K. Robinson, N. V. Roche, A. M. Smith, L. Sobin, D. E. Tabor, A. Undale, J. Bridge, L. E. Brigham, B. A. Foster, B. M. Gillard, R. Hasz, M. Hunter, C. Johns, M. Johnson, E. Karasik, G. Kopen, W. F. Leinweber, A. McDonald, M. T. Moser, K. Myer, K. D. Ramsey, B. Roe, S. Shad, J. A. Thomas, G. Walters, M. Washington, J. Wheeler, S. D. Jewell, D. C. Rohrer, D. R. Valley, D. A. Davis, D. C. Mash, P. A. Branton, L. K. Barker, H. M. Gardiner, M. Mosavel, L. A. Siminoff, P. Flicek, M. Haeussler, T. Juettemann, W. J. Kent, C. M. Lee, C. C. Powell, K. R. Rosenbloom, M. Ruffier, D. Sheppard, K. Taylor, S. J. Trevanion, D. R. Zerbino, N. S. Abell, J. Akey, L. Chen, K. Demanelis, J. A. Doherty, A. P. Feinberg, K. D. Hansen, P. F. Hickey, F. Jasmine, L. Jiang, R. Kaul, M. G. Kibriya, J. B. Li, Q. Li, S. Lin, S. E. Linder, B. L. Pierce, L. F. Rizzardi, A. D. Skol, K. S. Smith, M. Snyder, J. Stamatoyannopoulos, H. Tang, M. Wang, L. J. Carithers, P. Guan, S. E. Koester, A. R. Little, H. M. Moore, C. R. Nierras, A. K. Rao, J. B. Vaught, S. Volpi, The GTEx Consortium atlas of genetic regulatory effects across human tissues. Science 369, 1318–1330 (2020).

8. F. Aguet, A. A. Brown, S. E. Castel, J. R. Davis, Y. He, B. Jo, P. Mohammadi, Y. Park, P. Parsana, A. V. Segrè, B. J. Strober, Z. Zappala, B. B. Cummings, E. T. Gelfand, K. Hadley, K. H. Huang, M. Lek, X. Li, J. L. Nedzel, D. Y. Nguyen, M. S. Noble, T. J. Sullivan, T. Tukiainen, D. G. MacArthur, G. Getz, A. Addington, P. Guan, S. Koester, A. R. Little, N. C. Lockhart, H. M. Moore, A. Rao, J. P. Struewing, S. Volpi, L. E. Brigham, R. Hasz, M. Hunter, C. Johns, M. Johnson, G. Kopen, W. F. Leinweber, J. T. Lonsdale, A. McDonald, B. Mestichelli, K. Myer, B. Roe, M. Salvatore, S. Shad, J. A. Thomas, G. Walters, M. Washington, J. Wheeler, J. Bridge, B. A. Foster, B. M. Gillard, E. Karasik, R. Kumar, M. Miklos, M. T. Moser, S. D. Jewell, R. G. Montroy, D. C. Rohrer, D. Valley, D. C. Mash, D. A. Davis, L. Sobin, M. E. Barcus, P. A. Branton, N. S. Abell, B. Balliu, O. Delaneau, L. Frésard, E. R. Gamazon, D. Garrido-Martín, A. D. H. Gewirtz, G. Gliner, M. J. Gloudemans, B. Han, A. Z. He, F. Hormozdiari, X. Li, B. Liu, E. Y. Kang, I. C. McDowell, H. Ongen, J. J. Palowitch, C. B. Peterson, G. Quon, S. Ripke, A. Saha, A. A. Shabalin, T. C. Shimko, J. H. Sul, N. A. Teran, E. K. Tsang, H. Zhang, Y.-H. Zhou, C. D. Bustamante, N. J. Cox, R. Guigó, M. Kellis, M. I. McCarthy, D. F. Conrad, E. Eskin, G. Li, A. B. Nobel, C. Sabatti, B. E. Stranger, X. Wen, F. A. Wright, K. G. Ardlie, E. T. Dermitzakis, T. Lappalainen, F. Aguet, K. G. Ardlie, B. B. Cummings, E. T. Gelfand, G. Getz, K. Hadley, R. E. Handsaker, K. H. Huang, S. Kashin, K. J. Karczewski, M. Lek, X. Li, D. G. MacArthur, J. L. Nedzel, D. T. Nguyen, M. S. Noble, A. V. Segrè, C. A. Trowbridge, T. Tukiainen, N. S. Abell, B. Balliu, R. Barshir, O. Basha, A. Battle, G. K. Bogu, A. Brown, C. D. Brown, S. E. Castel, L. S. Chen, C. Chiang, D. F. Conrad, N. J. Cox, F. N. Damani, J. R. Davis, O. Delaneau, E. T. Dermitzakis, B. E. Engelhardt, E. Eskin, P. G. Ferreira, L. Frésard, E. R. Gamazon, D. Garrido-Martín, A. D. H. Gewirtz, G. Gliner, M. J. Gloudemans, R. Guigo, I. M. Hall, B. Han, Y. He, F. Hormozdiari, C. Howald, H. Kyung Im, B. Jo, E. Yong Kang, Y. Kim, S. Kim-Hellmuth, T. Lappalainen, G. Li, X. Li, B. Liu, S. Mangul, M. I. McCarthy, I. C. McDowell, P. Mohammadi, J. Monlong, S. B. Montgomery, M. Muñoz-Aguirre, A. W. Ndungu, D. L. Nicolae, A. B. Nobel, M. Oliva, H. Ongen, J. J. Palowitch, N. Panousis, P. Papasaikas, Y. Park, P. Parsana, A. J. Payne, C. B. Peterson, J. Quan, F. Reverter, C. Sabatti, A. Saha, M. Sammeth, A. J. Scott, A. A. Shabalin, R. Sodaei, M. Stephens, B. E. Stranger, B. J. Strober, J. H. Sul, E. K. Tsang, S. Urbut, M. van de Bunt, G. Wang, X. Wen, F. A. Wright, H. S. Xi, E. Yeger-Lotem, Z. Zappala, J. B. Zaugg, Y.-H. Zhou, J. M. Akey, D. Bates, J. Chan, L. S. Chen, M. Claussnitzer, K. Demanelis, M. Diegel, J. A. Doherty, A. P. Feinberg, M. S. Fernando, J. Halow, K. D. Hansen, E. Haugen, P. F. Hickey, L. Hou, F. Jasmine, R. Jian, L. Jiang, A. Johnson, R. Kaul, M. Kellis, M. G. Kibriya, K. Lee, J. Billy Li, Q. Li, X. Li, J. Lin, S. Lin, S. Linder, C. Linke, Y. Liu, M. T. Maurano, B. Molinie, S. B. Montgomery, J. Nelson, F. J. Neri, M. Oliva, Y. Park, B. L. Pierce, N. J. Rinaldi, L. F. Rizzardi, R. Sandstrom, A. Skol, K. S. Smith, M. P. Snyder, J. Stamatoyannopoulos, B. E. Stranger, H. Tang, E. K. Tsang, L. Wang, M. Wang, N. Van Wittenberghe, F. Wu, R. Zhang, C. R. Nierras, P. A. Branton, L. J. Carithers, P. Guan, H. M. Moore, A. Rao, J. B. Vaught, S. E. Gould, N. C. Lockart, C. Martin, J. P. Struewing, S. Volpi, A. M. Addington, S. E. Koester, A. R. Little, G. T. Consortium, a. Lead, D. A. Laboratory, C. Coordinating, N. I. H. p. management, c. Biospecimen, Pathology, Q. T. L. m. w. g. e, D. A. Laboratory, G. Coordinating Center —Analysis Working, G. Statistical Methods groups—Analysis Working, G. g. Enhancing, N. I. H. C. Fund, Nih/Nci, Nih/Nhgri, Nih/Nimh, Nih/Nida, N. Biospecimen Collection Source Site—, Genetic effects on gene expression across human tissues. Nature 550, 204–213 (2017).

9. A. S. E. Cuomo, A. Nathan, S. Raychaudhuri, D. G. MacArthur, J. E. Powell, Single-cell genomics meets human genetics. Nat Rev Genet, (2023).

10. H. Mostafavi, J. P. Spence, S. Naqvi, J. K. Pritchard, Limited overlap of eQTLs and GWAS hits due to systematic differences in discovery. bioRxiv, 2022.2005.2007.491045 (2022).

11. N. J. Connally, S. Nazeen, D. Lee, H. Shi, J. Stamatoyannopoulos, S. Chun, C. Cotsapas, C. A. Cassa, S. R. Sunyaev, The missing link between genetic association and regulatory function. eLife 11, e74970 (2022).

12. B. D. Umans, A. Battle, Y. Gilad, Where Are the Disease-Associated eQTLs? Trends Genet 37, 109–124 (2021).

13. K. A. Jagadeesh, K. K. Dey, D. T. Montoro, R. Mohan, S. Gazal, J. M. Engreitz, R. J. Xavier, A. L. Price, A. Regev, Identifying disease-critical cell types and cellular processes by integrating single-cell RNA-sequencing and human genetics. Nat. Genet. 54, 1479–1492 (2022).

14. G. Eraslan, E. Drokhlyansky, S. Anand, E. Fiskin, A. Subramanian, M. Slyper, J. Wang, N. Van Wittenberghe, J. M. Rouhana, J. Waldman, O. Ashenberg, M. Lek, D. Dionne, T. S. Win, M. S. Cuoco, O. Kuksenko, A. M. Tsankov, P. A. Branton, J. L. Marshall, A. Greka, G. Getz, A. V. Segre, F. Aguet, O. Rozenblatt-Rosen, K. G. Ardlie, A. Regev, Single-nucleus cross-tissue molecular reference maps toward understanding disease gene function. Science 376, eabl4290 (2022).

15. K. Zhang, J. D. Hocker, M. Miller, X. Hou, J. Chiou, O. B. Poirion, Y. Qiu, Y. E. Li, K. J. Gaulton, A. Wang, S. Preissl, B. Ren, A single-cell atlas of chromatin accessibility in the human genome. Cell 184, 5985–6001 e5919 (2021).

16. H. K. Finucane, Y. A. Reshef, V. Anttila, K. Slowikowski, A. Gusev, A. Byrnes, S. Gazal, P. R. Loh, C. Lareau, N. Shoresh, G. Genovese, A. Saunders, E. Macosko, S. Pollack, C. Brainstorm, J. R. B. Perry, J. D. Buenrostro, B. E. Bernstein, S. Raychaudhuri, S. McCarroll, B. M. Neale, A. L. Price, Heritability enrichment of specifically expressed genes identifies disease-relevant tissues and cell types. Nat. Genet. 50, 621–629 (2018).

17. X. Sheng, Y. Guan, Z. Ma, J. Wu, H. Liu, C. Qiu, S. Vitale, Z. Miao, M. J. Seasock, M. Palmer, M. K. Shin, K. L. Duffin, S. S. Pullen, T. L. Edwards, J. N. Hellwege, A. M. Hung, M. Li, B. F. Voight, T. M. Coffman, C. D. Brown, K. Susztak, Mapping the genetic architecture of human traits to cell types in the kidney identifies mechanisms of disease and potential treatments. Nat. Genet. 53, 1322–1333 (2021).

18. J. K. Lunney, A. Van Goor, K. E. Walker, T. Hailstock, J. Franklin, C. Dai, Importance of the pig as a human biomedical model. Sci. Transl. Med. 13, eabd5758 (2021).

19. Z.-L. Hu, C. A. Park, J. M. Reecy, Bringing the Animal QTLdb and CorrDB into the future: meeting new challenges and providing updated services. Nucleic Acids Research 50, D956–D961 (2021).

20. M. A. Groenen, A decade of pig genome sequencing: a window on pig domestication and evolution. Genet. Sel. Evol. 48, 23 (2016).

21. S. Liu, Y. Gao, O. Canela-Xandri, S. Wang, Y. Yu, W. Cai, B. Li, R. Xiang, A. J. Chamberlain, E. Pairo-Castineira, K. D’Mellow, K. Rawlik, C. Xia, Y. Yao, P. Navarro, D. Rocha, X. Li, Z. Yan, C. Li, B. D. Rosen, C. P. Van Tassell, P. M. Vanraden, S. Zhang, L. Ma, J. B. Cole, G. E. Liu, A. Tenesa, L. Fang, A multi-tissue atlas of regulatory variants in cattle. Nat. Genet. 54, 1438–1447 (2022).

22. Z. Pan, Y. Yao, H. Yin, Z. Cai, Y. Wang, L. Bai, C. Kern, M. Halstead, G. Chanthavixay, N. Trakooljul, K. Wimmers, G. Sahana, G. Su, M. S. Lund, M. Fredholm, P. Karlskov-Mortensen, C. W. Ernst, P. Ross, C. K. Tuggle, L. Fang, H. Zhou, Pig genome functional annotation enhances the biological interpretation of complex traits and human disease. Nat Commun 12, 5848 (2021).

23. L. Andersson, A. L. Archibald, C. D. Bottema, R. Brauning, S. C. Burgess, D. W. Burt, E. Casas, H. H. Cheng, L. Clarke, C. Couldrey, B. P. Dalrymple, C. G. Elsik, S. Foissac, E. Giuffra, M. A. Groenen, B. J. Hayes, L. S. Huang, H. Khatib, J. W. Kijas, H. Kim, J. K. Lunney, F. M. McCarthy, J. C. McEwan, S. Moore, B. Nanduri, C. Notredame, Y. Palti, G. S. Plastow, J. M. Reecy, G. A. Rohrer, E. Sarropoulou, C. J. Schmidt, J. Silverstein, R. L. Tellam, M. Tixier-Boichard, G. Tosser-Klopp, C. K. Tuggle, J. Vilkki, S. N. White, S. Zhao, H. Zhou, F. Consortium, Coordinated international action to accelerate genome-to-phenome with FAANG, the Functional Annotation of Animal Genomes project. Genome Biol 16, 57 (2015).

24. J. Teng, Y. Gao, H. Yin, Z. Bai, S. Liu, H. Zeng, L. Bai, Z. Cai, B. Zhao, X. Li, Z. Xu, Q. Lin, Z. Pan, W. Yang, X. Yu, D. Guan, Y. Hou, B. N. Keel, G. A. Rohrer, A. K. Lindholm-Perry, W. T. Oliver, M. Ballester, D. Crespo-Piazuelo, R. Quintanilla, O. Canela-Xandri, K. Rawlik, C. Xia, Y. Yao, Q. Zhao, W. Yao, L. Yang, H. Li, H. Zhang, W. Liao, T. Chen, P. Karlskov-Mortensen, M. Fredholm, M. Amills, A. Clop, E. Giuffra, J. Wu, X. Cai, S. Diao, X. Pan, C. Wei, J. Li, H. Cheng, S. Wang, G. Su, G. Sahana, M. S. Lund, J. C. M. Dekkers, L. Kramer, C. K. Tuggle, R. Corbett, M. A. M. Groenen, O. Madsen, M. Gòdia, D. Rocha, M. Charles, C.-j. Li, H. Pausch, X. Hu, L. Frantz, Y. Luo, L. Lin, Z. Zhou, Z. Zhang, Z. Chen, L. Cui, R. Xiang, X. Shen, P. Li, R. Huang, G. Tang, M. Li, Y. Zhao, G. Yi, Z. Tang, J. Jiang, F. Zhao, X. Yuan, X. Liu, Y. Chen, X. Xu, S. Zhao, P. Zhao, C. Haley, H. Zhou, Q. Wang, Y. Pan, X. Ding, L. Ma, J. Li, P. Navarro, Q. Zhang, B. Li, A. Tenesa, K. Li, G. E. Liu, Z. Zhang, L. Fang, A compendium of genetic regulatory effects across pig tissues. bioRxiv, 2022.2011.2011.516073 (2022).

25. L. Zhang, M. Guo, Z. Liu, R. Liu, Y. Zheng, T. Yu, Y. Lv, H. Lu, W. Zeng, T. Zhang, C. Pan, Single-cell RNA-seq analysis of testicular somatic cell development in pigs. J Genet Genomics 49, 1016–1028 (2022).

26. J. E. Wiarda, J. M. Trachsel, S. K. Sivasankaran, C. K. Tuggle, C. L. Loving, Intestinal single-cell atlas reveals novel lymphocytes in pigs with similarities to human cells. Life Science Alliance 5, e202201442 (2022).

27. F. Wang, P. Ding, X. Liang, X. Ding, C. B. Brandt, E. Sjöstedt, J. Zhu, S. Bolund, L. Zhang, L. P. M. H. de Rooij, L. Luo, Y. Wei, W. Zhao, Z. Lv, J. Haskó, R. Li, Q. Qin, Y. Jia, W. Wu, Y. Yuan, M. Pu, H. Wang, A. Wu, L. Xie, P. Liu, F. Chen, J. Herold, J. Kalucka, M. Karlsson, X. Zhang, R. B. Helmig, L. Fagerberg, C. Lindskog, F. Pontén, M. Uhlen, L. Bolund, N. Jessen, H. Jiang, X. Xu, H. Yang, P. Carmeliet, J. Mulder, D. Chen, L. Lin, Y. Luo, Endothelial cell heterogeneity and microglia regulons revealed by a pig cell landscape at single-cell level. Nature Communications 13, 3620 (2022).

28. L. Han, C. P. Jara, O. Wang, Y. Shi, X. Wu, S. Thibivilliers, R. K. Wóycicki, M. A. Carlson, W. H. Velander, E. P. Araújo, M. Libault, C. Zhang, Y. Lei, Isolating and cryopreserving pig skin cells for single-cell RNA sequencing study. PLoS One 17, e0263869 (2022).

29. J. Zhu, F. Chen, L. Luo, W. Wu, J. Dai, J. Zhong, X. Lin, C. Chai, P. Ding, L. Liang, S. Wang, X. Ding, Y. Chen, H. Wang, J. Qiu, F. Wang, C. Sun, Y. Zeng, J. Fang, X. Jiang, P. Liu, G. Tang, X. Qiu, X. Zhang, Y. Ruan, S. Jiang, J. Li, S. Zhu, X. Xu, F. Li, Z. Liu, G. Cao, D. Chen, Single-cell atlas of domestic pig cerebral cortex and hypothalamus. Science Bulletin 66, 1448–1461 (2021).

30. L. Zhang, J. Zhu, H. Wang, J. Xia, P. Liu, F. Chen, H. Jiang, Q. Miao, W. Wu, L. Zhang, L. Luo, X. Jiang, Y. Bai, C. Sun, D. Chen, X. Zhang, A high-resolution cell atlas of the domestic pig lung and an online platform for exploring lung single-cell data. J Genet Genomics 48, 411–425 (2021).

31. J. Herrera-Uribe, J. E. Wiarda, S. K. Sivasankaran, L. Daharsh, H. Liu, K. A. Byrne, T. P. L. Smith, J. K. Lunney, C. L. Loving, C. K. Tuggle, Reference Transcriptomes of Porcine Peripheral Immune Cells Created Through Bulk and Single-Cell RNA Sequencing. Front. Genet. 12, (2021).

32. W. Tang, Y. Zhong, Y. Wei, Z. Deng, J. Mao, J. Liu, T. G. Valencak, J. Liu, H. Xu, H. Wang, Ileum tissue single-cell mRNA sequencing elucidates the cellular architecture of pathophysiological changes associated with weaning in piglets. BMC Biology 20, 123 (2022).

33. H. L. Sweeney, D. W. Hammers, Muscle Contraction. Cold Spring Harb. Perspect. Biol. 10, (2018).

34. K. Mukund, S. Subramaniam, Skeletal muscle: A review of molecular structure and function, in health and disease. Wiley Interdiscip. Rev. Syst. Biol. Med. 12, e1462 (2020).

35. P. Orchard, N. Manickam, C. Ventresca, S. Vadlamudi, A. Varshney, V. Rai, J. Kaplan, C. Lalancette, K. L. Mohlke, K. Gallagher, C. F. Burant, S. C. J. Parker, Human and rat skeletal muscle single-nuclei multi-omic integrative analyses nominate causal cell types, regulatory elements, and SNPs for complex traits. Genome Res. 31, 2258–2275 (2021).

36. A. B. Rubenstein, G. R. Smith, U. Raue, G. Begue, K. Minchev, F. Ruf-Zamojski, V. D. Nair, X. Wang, L. Zhou, E. Zaslavsky, T. A. Trappe, S. Trappe, S. C. Sealfon, Single-cell transcriptional profiles in human skeletal muscle. Sci. Rep. 10, 229 (2020).

37. J. Günther, H.-M. Seyfert, The first line of defence: insights into mechanisms and relevance of phagocytosis in epithelial cells. Semin. Immunopathol. 40, 555–565 (2018).

38. R. Elmentaite, N. Kumasaka, K. Roberts, A. Fleming, E. Dann, H. W. King, V. Kleshchevnikov, M. Dabrowska, S. Pritchard, L. Bolt, S. F. Vieira, L. Mamanova, N. Huang, F. Perrone, I. Goh Kai’En, S. N. Lisgo, M. Katan, S. Leonard, T. R. W. Oliver, C. E. Hook, K. Nayak, L. S. Campos, C. Dominguez Conde, E. Stephenson, J. Engelbert, R. A. Botting, K. Polanski, S. van Dongen, M. Patel, M. D. Morgan, J. C. Marioni, O. A. Bayraktar, K. B. Meyer, X. He, R. A. Barker, H. H. Uhlig, K. T. Mahbubani, K. Saeb-Parsy, M. Zilbauer, M. R. Clatworthy, M. Haniffa, K. R. James, S. A. Teichmann, Cells of the human intestinal tract mapped across space and time. Nature 597, 250–255 (2021).

39. A. Ali, H. Tan, G. E. Kaiko, Role of the Intestinal Epithelium and Its Interaction With the Microbiota in Food Allergy. Front. Immunol. 11, 604054 (2020).

40. A. L. Haber, M. Biton, N. Rogel, R. H. Herbst, K. Shekhar, C. Smillie, G. Burgin, T. M. Delorey, M. R. Howitt, Y. Katz, I. Tirosh, S. Beyaz, D. Dionne, M. Zhang, R. Raychowdhury, W. S. Garrett, O. Rozenblatt-Rosen, H. N. Shi, O. Yilmaz, R. J. Xavier, A. Regev, A single-cell survey of the small intestinal epithelium. Nature 551, 333–339 (2017).

41. J. Cao, M. Spielmann, X. Qiu, X. Huang, D. M. Ibrahim, A. J. Hill, F. Zhang, S. Mundlos, L. Christiansen, F. J. Steemers, C. Trapnell, J. Shendure, The single-cell transcriptional landscape of mammalian organogenesis. Nature 566, 496–502 (2019).

42. S. Jin, C. F. Guerrero-Juarez, L. Zhang, I. Chang, R. Ramos, C. H. Kuan, P. Myung, M. V. Plikus, Q. Nie, Inference and analysis of cell-cell communication using CellChat. Nat Commun 12, 1088 (2021).

43. D. D. Chaplin, Overview of the immune response. J Allergy Clin Immunol 125, S3–23 (2010).

44. A. M. Newman, C. L. Liu, M. R. Green, A. J. Gentles, W. Feng, Y. Xu, C. D. Hoang, M. Diehn, A. A. Alizadeh, Robust enumeration of cell subsets from tissue expression profiles. Nat. Methods 12, 453–457 (2015).

45. M. Dong, A. Thennavan, E. Urrutia, Y. Li, C. M. Perou, F. Zou, Y. Jiang, SCDC: bulk gene expression deconvolution by multiple single-cell RNA sequencing references. Brief Bioinform 22, 416–427 (2021).

46. S. Kim-Hellmuth, F. Aguet, M. Oliva, M. Muñoz-Aguirre, S. Kasela, V. Wucher, S. E. Castel, A. R. Hamel, A. Viñuela, A. L. Roberts, S. Mangul, X. Wen, G. Wang, A. N. Barbeira, D. Garrido-Martín, B. B. Nadel, Y. Zou, R. Bonazzola, J. Quan, A. Brown, A. Martinez-Perez, J. M. Soria, G. Getz, E. T. Dermitzakis, K. S. Small, M. Stephens, H. S. Xi, H. K. Im, R. Guigó, A. V. Segrè, B. E. Stranger, K. G. Ardlie, T. Lappalainen, Cell type-specific genetic regulation of gene expression across human tissues. Science 369, (2020).

47. Y. Yamazaki, N. Zhao, T. R. Caulfield, C. C. Liu, G. Bu, Apolipoprotein E and Alzheimer disease: pathobiology and targeting strategies. Nat. Rev. Neurol. 15, 501–518 (2019).

48. H. Van Gorp, W. Van Breedam, J. Van Doorsselaere, P. L. Delputte, H. J. Nauwynck, Identification of the CD163 protein domains involved in infection of the porcine reproductive and respiratory syndrome virus. J. Virol. 84, 3101–3105 (2010).

49. J. J. Wu, S. Zhu, F. Gu, T. G. Valencak, J. X. Liu, H. Z. Sun, Cross-tissue single-cell transcriptomic landscape reveals the key cell subtypes and their potential roles in the nutrient absorption and metabolism in dairy cattle. J Adv Res 37, 1–18 (2022).

50. J. Qu, F. Yang, T. Zhu, Y. Wang, W. Fang, Y. Ding, X. Zhao, X. Qi, Q. Xie, M. Chen, Q. Xu, Y. Xie, Y. Sun, D. Chen, A reference single-cell regulomic and transcriptomic map of cynomolgus monkeys. Nat Commun 13, 4069 (2022).

51. H. Li, J. Janssens, M. De Waegeneer, S. S. Kolluru, K. Davie, V. Gardeux, W. Saelens, F. P. A. David, M. Brbić, K. Spanier, J. Leskovec, C. N. McLaughlin, Q. Xie, R. C. Jones, K. Brueckner, J. Shim, S. G. Tattikota, F. Schnorrer, K. Rust, T. G. Nystul, Z. Carvalho-Santos, C. Ribeiro, S. Pal, S. Mahadevaraju, T. M. Przytycka, A. M. Allen, S. F. Goodwin, C. W. Berry, M. T. Fuller, H. White-Cooper, E. L. Matunis, S. DiNardo, A. Galenza, L. E. O’Brien, J. A. T. Dow, H. Jasper, B. Oliver, N. Perrimon, B. Deplancke, S. R. Quake, L. Luo, S. Aerts, D. Agarwal, Y. Ahmed-Braimah, M. Arbeitman, M. M. Ariss, J. Augsburger, K. Ayush, C. C. Baker, T. Banisch, K. Birker, R. Bodmer, B. Bolival, S. E. Brantley, J. A. Brill, N. C. Brown, N. A. Buehner, X. T. Cai, R. Cardoso-Figueiredo, F. Casares, A. Chang, T. R. Clandinin, S. Crasta, C. Desplan, A. M. Detweiler, D. B. Dhakan, E. Donà, S. Engert, S. Floc’hlay, N. George, A. J. González-Segarra, A. K. Groves, S. Gumbin, Y. Guo, D. E. Harris, Y. Heifetz, S. L. Holtz, F. Horns, B. Hudry, R. J. Hung, Y. N. Jan, J. S. Jaszczak, G. Jefferis, J. Karkanias, T. L. Karr, N. S. Katheder, J. Kezos, A. A. Kim, S. K. Kim, L. Kockel, N. Konstantinides, T. B. Kornberg, H. M. Krause, A. T. Labott, M. Laturney, R. Lehmann, S. Leinwand, J. Li, J. S. S. Li, K. Li, K. Li, L. Li, T. Li, M. Litovchenko, H. H. Liu, Y. Liu, T. C. Lu, J. Manning, A. Mase, M. Matera-Vatnick, N. R. Matias, C. E. McDonough-Goldstein, A. McGeever, A. D. McLachlan, P. Moreno-Roman, N. Neff, M. Neville, S. Ngo, T. Nielsen, C. E. O’Brien, D. Osumi-Sutherland, M. N. Özel, I. Papatheodorou, M. Petkovic, C. Pilgrim, A. O. Pisco, C. Reisenman, E. N. Sanders, G. Dos Santos, K. Scott, A. Sherlekar, P. Shiu, D. Sims, R. V. Sit, M. Slaidina, H. E. Smith, G. Sterne, Y. H. Su, D. Sutton, M. Tamayo, M. Tan, I. Tastekin, C. Treiber, D. Vacek, G. Vogler, S. Waddell, W. Wang, R. I. Wilson, M. F. Wolfner, Y. E. Wong, A. Xie, J. Xu, S. Yamamoto, J. Yan, Z. Yao, K. Yoda, R. Zhu, R. P. Zinzen, Fly Cell Atlas: A single-nucleus transcriptomic atlas of the adult fruit fly. Science 375, eabk2432 (2022).

52. J. Li, J. Wang, P. Zhang, R. Wang, Y. Mei, Z. Sun, L. Fei, M. Jiang, L. Ma, W. E, H. Chen, X. Wang, Y. Fu, H. Wu, D. Liu, X. Wang, J. Li, Q. Guo, Y. Liao, C. Yu, D. Jia, J. Wu, S. He, H. Liu, J. Ma, K. Lei, J. Chen, X. Han, G. Guo, Deep learning of cross-species single-cell landscapes identifies conserved regulatory programs underlying cell types. Nat. Genet. 54, 1711–1720 (2022).

53. Y. M. Choi, B. C. Kim, Muscle fiber characteristics, myofibrillar protein isoforms, and meat quality. Livestock Science 122, 105–118 (2009).

54. R. E. Klont, L. Brocks, G. Eikelenboom, Muscle fibre type and meat quality. Meat Sci 49s1, S219–229 (1998).

55. L. Jin, Q. Tang, S. Hu, Z. Chen, X. Zhou, B. Zeng, Y. Wang, M. He, Y. Li, L. Gui, L. Shen, K. Long, J. Ma, X. Wang, Z. Chen, Y. Jiang, G. Tang, L. Zhu, F. Liu, B. Zhang, Z. Huang, G. Li, D. Li, V. N. Gladyshev, J. Yin, Y. Gu, X. Li, M. Li, A pig BodyMap transcriptome reveals diverse tissue physiologies and evolutionary dynamics of transcription. Nat Commun 12, 3715 (2021).

56. J. Talbot, L. Maves, Skeletal muscle fiber type: using insights from muscle developmental biology to dissect targets for susceptibility and resistance to muscle disease. Wiley Interdiscip Rev Dev Biol 5, 518–534 (2016).

57. S. T. Joo, G. D. Kim, Y. H. Hwang, Y. C. Ryu, Control of fresh meat quality through manipulation of muscle fiber characteristics. Meat Sci 95, 828–836 (2013).

58. C. Domínguez Conde, C. Xu, L. B. Jarvis, D. B. Rainbow, S. B. Wells, T. Gomes, S. K. Howlett, O. Suchanek, K. Polanski, H. W. King, L. Mamanova, N. Huang, P. A. Szabo, L. Richardson, L. Bolt, E. S. Fasouli, K. T. Mahbubani, M. Prete, L. Tuck, N. Richoz, Z. K. Tuong, L. Campos, H. S. Mousa, E. J. Needham, S. Pritchard, T. Li, R. Elmentaite, J. Park, E. Rahmani, D. Chen, D. K. Menon, O. A. Bayraktar, L. K. James, K. B. Meyer, N. Yosef, M. R. Clatworthy, P. A. Sims, D. L. Farber, K. Saeb-Parsy, J. L. Jones, S. A. Teichmann, Cross-tissue immune cell analysis reveals tissue-specific features in humans. Science 376, eabl5197 (2022).

59. M. Slyper, C. B. M. Porter, O. Ashenberg, J. Waldman, E. Drokhlyansky, I. Wakiro, C. Smillie, G. Smith-Rosario, J. Wu, D. Dionne, S. Vigneau, J. Jane-Valbuena, T. L. Tickle, S. Napolitano, M. J. Su, A. G. Patel, A. Karlstrom, S. Gritsch, M. Nomura, A. Waghray, S. H. Gohil, A. M. Tsankov, L. Jerby-Arnon, O. Cohen, J. Klughammer, Y. Rosen, J. Gould, L. Nguyen, M. Hofree, P. J. Tramontozzi, B. Li, C. J. Wu, B. Izar, R. Haq, F. S. Hodi, C. H. Yoon, A. N. Hata, S. J. Baker, M. L. Suva, R. Bueno, E. H. Stover, M. R. Clay, M. A. Dyer, N. B. Collins, U. A. Matulonis, N. Wagle, B. E. Johnson, A. Rotem, O. Rozenblatt-Rosen, A. Regev, A single-cell and single-nucleus RNA-Seq toolbox for fresh and frozen human tumors. Nat. Med. 26, 792–802 (2020).

60. J. Ding, X. Adiconis, S. K. Simmons, M. S. Kowalczyk, C. C. Hession, N. D. Marjanovic, T. K. Hughes, M. H. Wadsworth, T. Burks, L. T. Nguyen, J. Y. H. Kwon, B. Barak, W. Ge, A. J. Kedaigle, S. Carroll, S. Li, N. Hacohen, O. Rozenblatt-Rosen, A. K. Shalek, A. C. Villani, A. Regev, J. Z. Levin, Systematic comparison of single-cell and single-nucleus RNA-sequencing methods. Nat. Biotechnol. 38, 737–746 (2020).

61. A. Warr, N. Affara, B. Aken, H. Beiki, D. M. Bickhart, K. Billis, W. Chow, L. Eory, H. A. Finlayson, P. Flicek, C. G. Giron, D. K. Griffin, R. Hall, G. Hannum, T. Hourlier, K. Howe, D. A. Hume, O. Izuogu, K. Kim, S. Koren, H. Liu, N. Manchanda, F. J. Martin, D. J. Nonneman, R. E. O’Connor, A. M. Phillippy, G. A. Rohrer, B. D. Rosen, L. A. Rund, C. A. Sargent, L. B. Schook, S. G. Schroeder, A. S. Schwartz, B. M. Skinner, R. Talbot, E. Tseng, C. K. Tuggle, M. Watson, T. P. L. Smith, A. L. Archibald, An improved pig reference genome sequence to enable pig genetics and genomics research. Gigascience 9, (2020).

62. R. Satija, J. A. Farrell, D. Gennert, A. F. Schier, A. Regev, Spatial reconstruction of single-cell gene expression data. Nat. Biotechnol. 33, 495–502 (2015).

63. S. Yang, S. E. Corbett, Y. Koga, Z. Wang, W. E. Johnson, M. Yajima, J. D. Campbell, Decontamination of ambient RNA in single-cell RNA-seq with DecontX. Genome Biol 21, 57 (2020).

64. C. S. McGinnis, L. M. Murrow, Z. J. Gartner, DoubletFinder: Doublet Detection in Single-Cell RNA Sequencing Data Using Artificial Nearest Neighbors. Cell Syst 8, 329–337.e324 (2019).

65. B. Hie, B. Bryson, B. Berger, Efficient integration of heterogeneous single-cell transcriptomes using Scanorama. Nat. Biotechnol. 37, 685–691 (2019).

66. V. Bergen, M. Lange, S. Peidli, F. A. Wolf, F. J. Theis, Generalizing RNA velocity to transient cell states through dynamical modeling. Nat. Biotechnol. 38, 1408–1414 (2020).

67. R. Xue, Q. Zhang, Q. Cao, R. Kong, X. Xiang, H. Liu, M. Feng, F. Wang, J. Cheng, Z. Li, Q. Zhan, M. Deng, J. Zhu, Z. Zhang, N. Zhang, Liver tumour immune microenvironment subtypes and neutrophil heterogeneity. Nature 612, 141–147 (2022).

68. T. Wu, E. Hu, S. Xu, M. Chen, P. Guo, Z. Dai, T. Feng, L. Zhou, W. Tang, L. Zhan, X. Fu, S. Liu, X. Bo, G. Yu, clusterProfiler 4.0: A universal enrichment tool for interpreting omics data. Innovation (Camb) 2, 100141 (2021).

69. S. Aibar, C. B. González-Blas, T. Moerman, V. A. Huynh-Thu, H. Imrichova, G. Hulselmans, F. Rambow, J. C. Marine, P. Geurts, J. Aerts, J. van den Oord, Z. K. Atak, J. Wouters, S. Aerts, SCENIC: single-cell regulatory network inference and clustering. Nat. Methods 14, 1083–1086 (2017).

70. H. Jin, Z. Liu, A benchmark for RNA-seq deconvolution analysis under dynamic testing environments. Genome Biol 22, 102 (2021).

71. A. Taylor-Weiner, F. Aguet, N. J. Haradhvala, S. Gosai, S. Anand, J. Kim, K. Ardlie, E. M. Van Allen, G. Getz, Scaling computational genomics to millions of individuals with GPUs. Genome Biol 20, 228 (2019).

72. J. R. Davis, L. Fresard, D. A. Knowles, M. Pala, C. D. Bustamante, A. Battle, S. B. Montgomery, An Efficient Multiple-Testing Adjustment for eQTL Studies that Accounts for Linkage Disequilibrium between Variants. Am. J. Hum. Genet. 98, 216–224 (2016).

73. S. E. Castel, P. Mohammadi, W. K. Chung, Y. Shen, T. Lappalainen, Rare variant phasing and haplotypic expression from RNA sequencing with phASER. Nat Commun 7, 12817 (2016).

74. S. E. Castel, A. Levy-Moonshine, P. Mohammadi, E. Banks, T. Lappalainen, Tools and best practices for data processing in allelic expression analysis. Genome Biol 16, 195 (2015).

75. C. Giambartolomei, D. Vukcevic, E. E. Schadt, L. Franke, A. D. Hingorani, C. Wallace, V. Plagnol, Bayesian test for colocalisation between pairs of genetic association studies using summary statistics. PLoS Genet 10, e1004383 (2014).

76. N. C. Sheffield, C. Bock, LOLA: enrichment analysis for genomic region sets and regulatory elements in R and Bioconductor. Bioinformatics 32, 587–589 (2016).

77. A. Marceau, Y. Gao, R. L. t. Baldwin, C. J. Li, J. Jiang, G. E. Liu, L. Ma, Investigation of rumen long noncoding RNA before and after weaning in cattle. BMC Genomics 23, 531 (2022).

78. H. K. Finucane, B. Bulik-Sullivan, A. Gusev, G. Trynka, Y. Reshef, P. R. Loh, V. Anttila, H. Xu, C. Zang, K. Farh, S. Ripke, F. R. Day, S. Purcell, E. Stahl, S. Lindstrom, J. R. Perry, Y. Okada, S. Raychaudhuri, M. J. Daly, N. Patterson, B. M. Neale, A. L. Price, Partitioning heritability by functional annotation using genome-wide association summary statistics. Nat. Genet. 47, 1228–1235 (2015).

79. B. K. Bulik-Sullivan, P. R. Loh, H. K. Finucane, S. Ripke, J. Yang, N. Patterson, M. J. Daly, A. L. Price, B. M. Neale, LD Score regression distinguishes confounding from polygenicity in genome-wide association studies. Nat. Genet. 47, 291–295 (2015).

